# Computational Assessment of Different Structural Models for Claudin-5 Complexes in Blood-Brain-Barrier Tight-Junctions

**DOI:** 10.1101/2022.03.03.482846

**Authors:** Alessandro Berselli, Giulio Alberini, Fabio Benfenati, Luca Maragliano

**Affiliations:** Center for Synaptic Neuroscience and Technology (NSYN@UniGe), Istituto Italiano di Tecnologia, Largo Rosanna Benzi, 10, 16132 Genova, Italy; Department of Experimental Medicine, Università degli Studi di Genova, Viale Benedetto XV, 3, 16132 Genova, Italy; IRCCS Ospedale Policlinico San Martino, Largo Rosanna Benzi, 10, 16132 Genova, Italy; Department of Life and Environmental Sciences, Polytechnic University of Marche, Via Brecce Bianche, 60131, Ancona, Italy

**Author notes:** Alessandro Berselli and Giulio Alberini equally contributed to perform the simulations, analyze the data, discuss the results and write the manuscript.

**Keywords:** Tight junctions, Claudin-5, Blood-brain barrier, Biological pore models, Molecular dynamics, Free energy calculations

## Abstract

The blood-brain barrier (BBB) strictly regulates the exchange of ions and molecules between the blood and the central nervous system. Tight junctions (TJs) are multimeric structures that control the transport through the paracellular spaces between adjacent brain endothelial cells of the BBB. Claudin-5 (Cldn5) proteins are essential for the TJ formation and assemble into multi-protein complexes via *cis*-interactions within the same cell membrane and *trans*-interactions across two contiguous cells. Despite the relevant biological function of Cldn5 proteins and their role as targets of brain drug delivery strategies, the molecular details of their assembly within TJs are still unclear. Two different structural models have been recently introduced, in which Cldn5 dimers belonging to opposite cells join to generate paracellular pores. However, a comparison of these models in terms of ionic transport features is still lacking. In this work, we used molecular dynamics simulations and free energy (FE) calculations to assess the two Cldn5 pore models and investigate the thermodynamic properties of water and physiological ions permeating through them. Despite different FE profiles, both structures present single/multiple FE barriers to ionic permeation, while being permissive to water flux. These results reveal that both models are compatible with the physiological role of Cldn5 TJ strands. By identifying the protein-protein surface at the core of TJ Cldn5 assemblies, our computational investigation provides a basis for the rational design of synthetic peptides and other molecules capable of opening paracellular pores in the BBB.

## INTRODUCTION

Biological barriers are structures made of layers of tightly bound endothelial/epithelial cells that preserve the characteristics of the body compartments they separate and regulate the exchanges between them. Multimeric protein complexes named *tight junctions* (TJs)^1–6^ hold adjacent cells together by forming strands that are visible in freeze-fracture electron microscopy images and seal the paracellular space between cells^7–9^.

Claudins (Cldns) are the major components of the TJ strands^7,10,11^. The Cldn family is composed of 27 tissuespecific homologs with a structure comprising a transmembrane four-helix bundle (TM1-4) embedded in the membrane bilayer (the TM domain). TM helices are joined by two extracellular loops spanning the paracellular space (ECL1-2) and by an intracellular loop in the cytoplasmic region, where the N/C termini are also found^12,13^. Cldns are known to assemble into TJs via intermolecular *cis*-interactions between individual protomers and *trans*-interactions between proteins of adjacent cells^14^. TJ strands regulate the paracellular flux of ions and molecules across the various barriers via highly selective, tissue-specific mechanisms^15,16^.

The Cldn subtype 5 (Cldn5) is the most abundant TJ protein in the endothelial cells of the blood-brain barrier (BBB), the highly selective interface that preserves the chemical homeostasis of the central nervous system (CNS). In particular, Cldn5 strands are responsible for the very limited BBB paracellular permeability that prevents the uncontrolled permeation of ions and small molecules^17–20^. The relevant physiological function of Cldn5 proteins makes them a novel and promising target for strategies to deliver drugs directly to the brain^21–25^. However, structure-based approaches are still hampered by a lack of knowledge on the precise assembly of Cldn5 protomers in the BBB TJs^17^. Only recently^25^, based on prior results of other Cldns^26–28^, two structural models of Cldn5 complexes were introduced, both of which display a pore cavity, and were named Pore I and Pore II.

The Pore I structure is based on the model originally introduced by Suzuki et al. in Ref. 29 for the homologous Cldn subtype 15 (Cldn15, PDB ID: 4P79)^30^, the first member of the family to be crystallized. According to this template, *cis*-interactions are formed by the ECL1 domains of two neighboring protomers in the same membrane (also named *face-to-face* interaction^7^), with opposing *β*-strands arranged in an antiparallel fashion to generate an extended *β*-sheet across the two molecules, defining a hydrophilic surface. Moreover, two opposing dimers from adjacent cells create a tetrameric arrangement sustained by *trans*-interactions between the ECLs of the protomers, resulting in a *β*-barrel super-secondary structure in the paracellular space that encompasses a pore cavity. After the publication of the Cldn15-based model, this has been refined and validated in several studies using structural modeling and molecular dynamics (MD) simulations^31–38^, also for other Cldns. In particular, in Ref. 37, the authors investigated the mechanism of ion permeation through the Cldn5 Pore I by calculating the free energy (FE, or potential of mean force, PMF) profiles for various ionic species. Results pointed to the lack of both cation and anion permeation, thus demonstrating that the Pore I conformation properly reproduces the function of barrier to ionic fluxes exerted by BBB TJs^39^.

On the other hand, the Pore II was also introduced by the same group^34,35,40^, based on previously modeled Cldn5 dimers^34^. Although the structure still comprises again two facing Cldn dimers, the *cis*-arrangement between two protomers in the membrane is characterized by a distinct pattern of interactions involving the TM2 and TM3 helices (also named *back-to-back* interaction^7^). More specifically, the authors identified a *leucine zipper* motif defined by the residues Leu83, Leu90, Leu124 and Leu131 of the two Cldn5 subunits, supported by the aromatic interactions between the opposing pairs of Trp138 and Phe127 residues. The presence of this *cis*-dimerization interface is consistent with the experimental results illustrated in Ref. 28. Then, similarly to the Pore I configuration^13,40^, the Pore II architecture is obtained by joining a couple of these dimers via *trans*-interactions, although it lacks the cavity-enveloping super-secondary structure of Pore I. The MD simulations illustrated in Ref. 40 demonstrate that the Cldn5 Pore II is impermeable to small molecules such as α-D glucose, but permissive to water. However, at variance with Pore I, the Pore II model is still limitedly investigated^34,35,40–42^, and further studies are required to chart its structural and functional hallmarks. Moreover, a detailed investigation of its ionic permeability has not been performed yet, thus hampering a thorough comparison with Pore I.

The aim of this work is to investigate the two different pore models and to assess their reliability as possible representatives of Cldn5 complexes in the BBB TJs. After building the two tetrameric configurations using Cldn5 protomers modeled from the homologous Cldn15^30^, we used all-atom MD simulations to refine their structures in solvated, double-membrane environments and to compute the one-dimensional FE profiles for the permeation of water and ions through both pores. Results show that the Pore I arrangement is structurally more stable, while both are water-permeable and present FE barriers of different heights to the passage of ions, consistently with the known role of Cldn5 in increasing the trans-endothelial electrical resistance and reducing the ionic paracellular permeability of the BBB^20^. In both the conformations, the FE critical points correlate with the positions of pore-lining charged residues. In particular, barriers for cations are localized in proximity of the Lys65 sidechains, while those for Cl^-^ are in correspondence of Glu146 and Asp149. The profiles for the same ions are, however, quite different in the two structures, due to distinct arrangements of the residues along the pores. Moreover, the hydration pattern of permeating ions along the pore axis shows a partial depletion of the coordinating water molecules in correspondence with the narrow regions of the pores^33,43^.

Our findings provide a systematic description of the two Cldn5 tetrameric pore configurations in terms of their structural and permeation properties, indicating that they are both possible Cldn5 assemblies in the TJs of the brain endothelium.

## RESULTS AND DISCUSSION

The tetrameric structures of Pore I and Pore II are shown in **Figures 1-3**, which report the arrangements of Cldn5 protomers in dimers (**Figure 1**), the quaternary structure of the two pores (**Figure 2**), and the relevant amino acids within their cavities (**Figure 3**).

**Figure 1.**
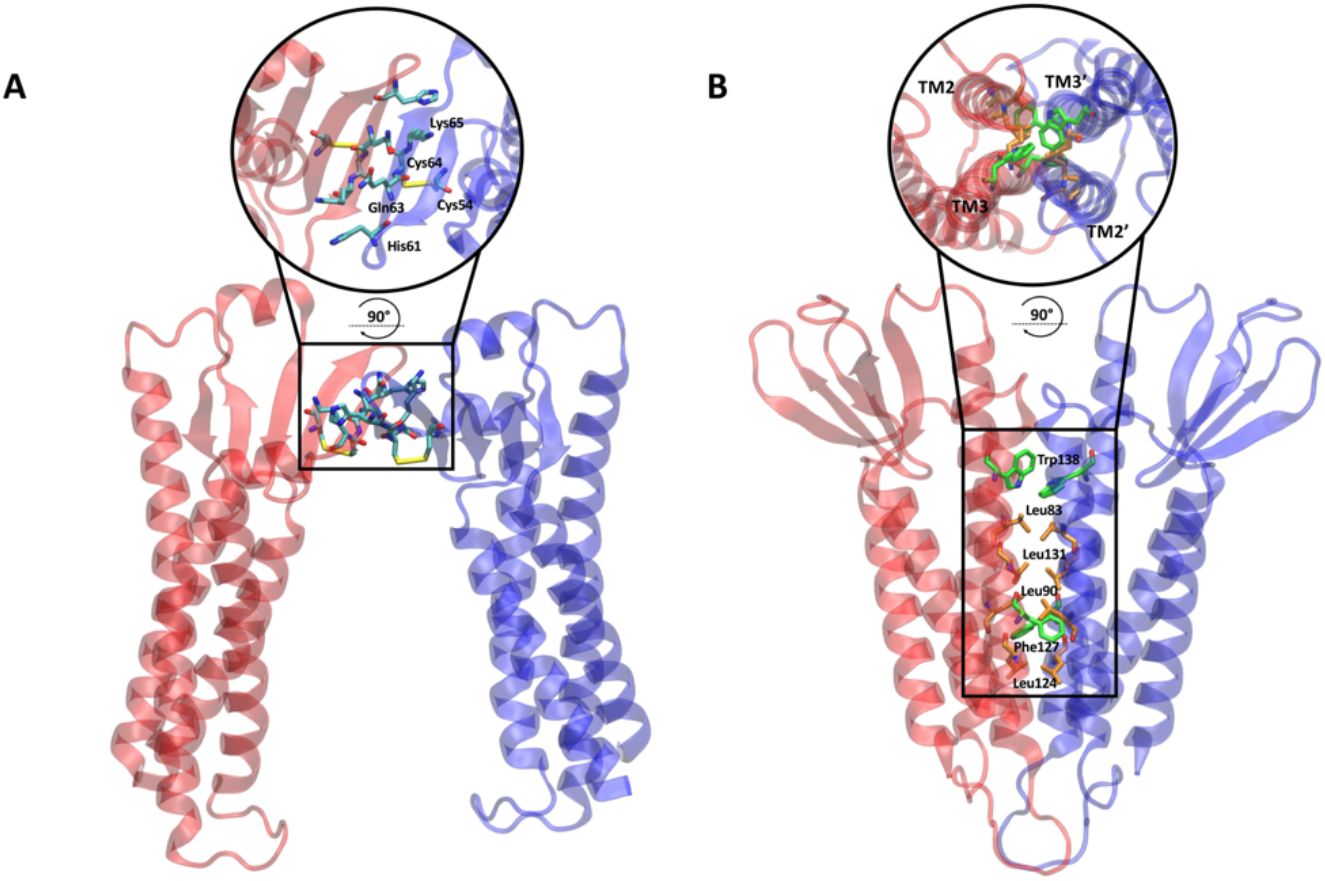
Structural representations of the two equilibrated dimeric structures which prelude to Pore I (A) and Pore II (B). The dimer in panel A is characterized by a face-to-face interaction between the ECL1 domains of two opposite Cldn5 protomers. The dimer in panel B is formed by a back-to-back interaction and stabilized by a leucine zipper pairs in the TM2-TM3 helices of the single Cldn5 protomers.

**Figure 2.**
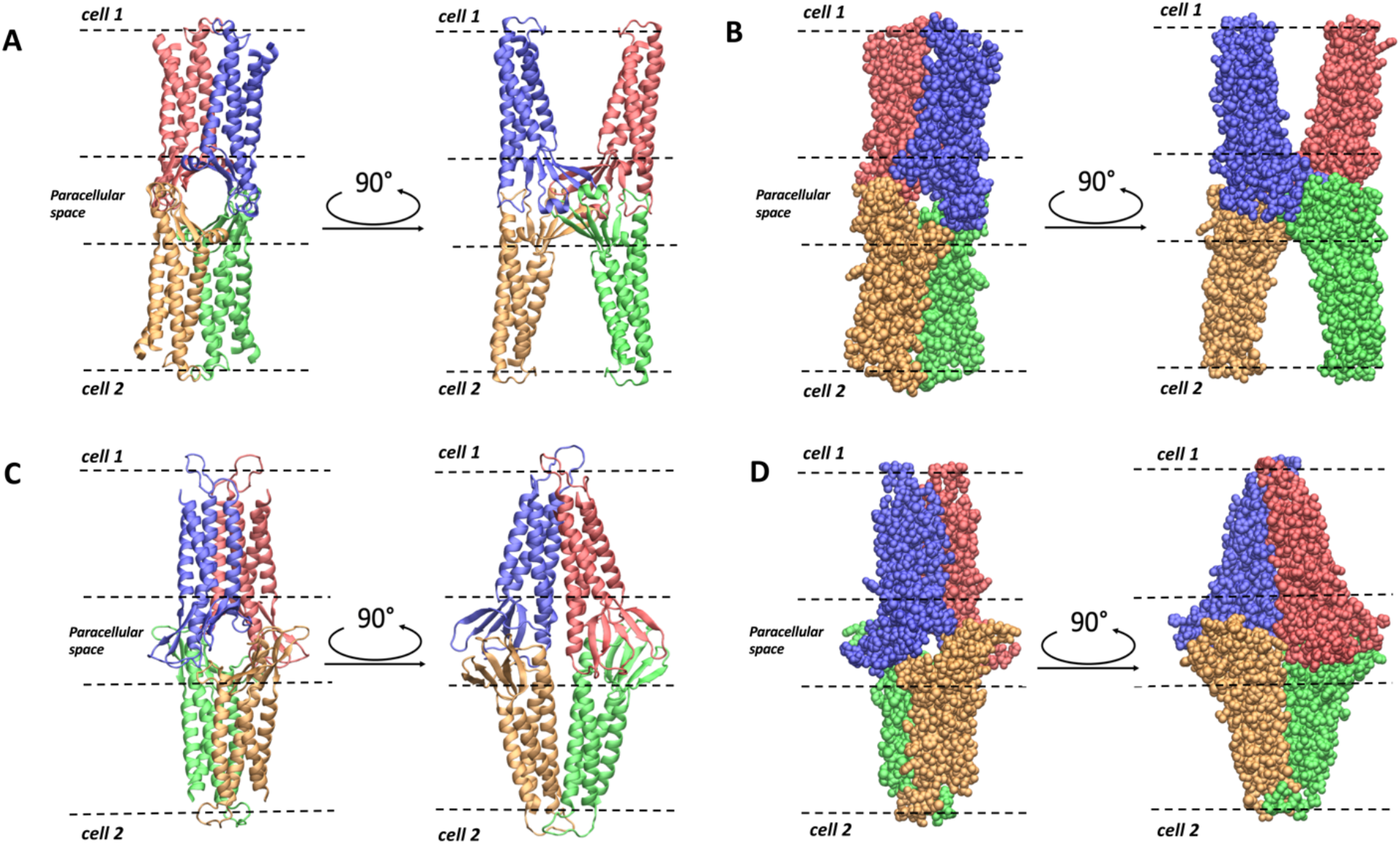
Structural representation of the two equilibrated single-pore models, top and side views. Pore configurations are shown in ribbon cartoon style (A, C) and Van Der Waals sphere style (B,D) for Pore I (A,B) and Pore II (C,D) models, respectively.

**Figure 3.**
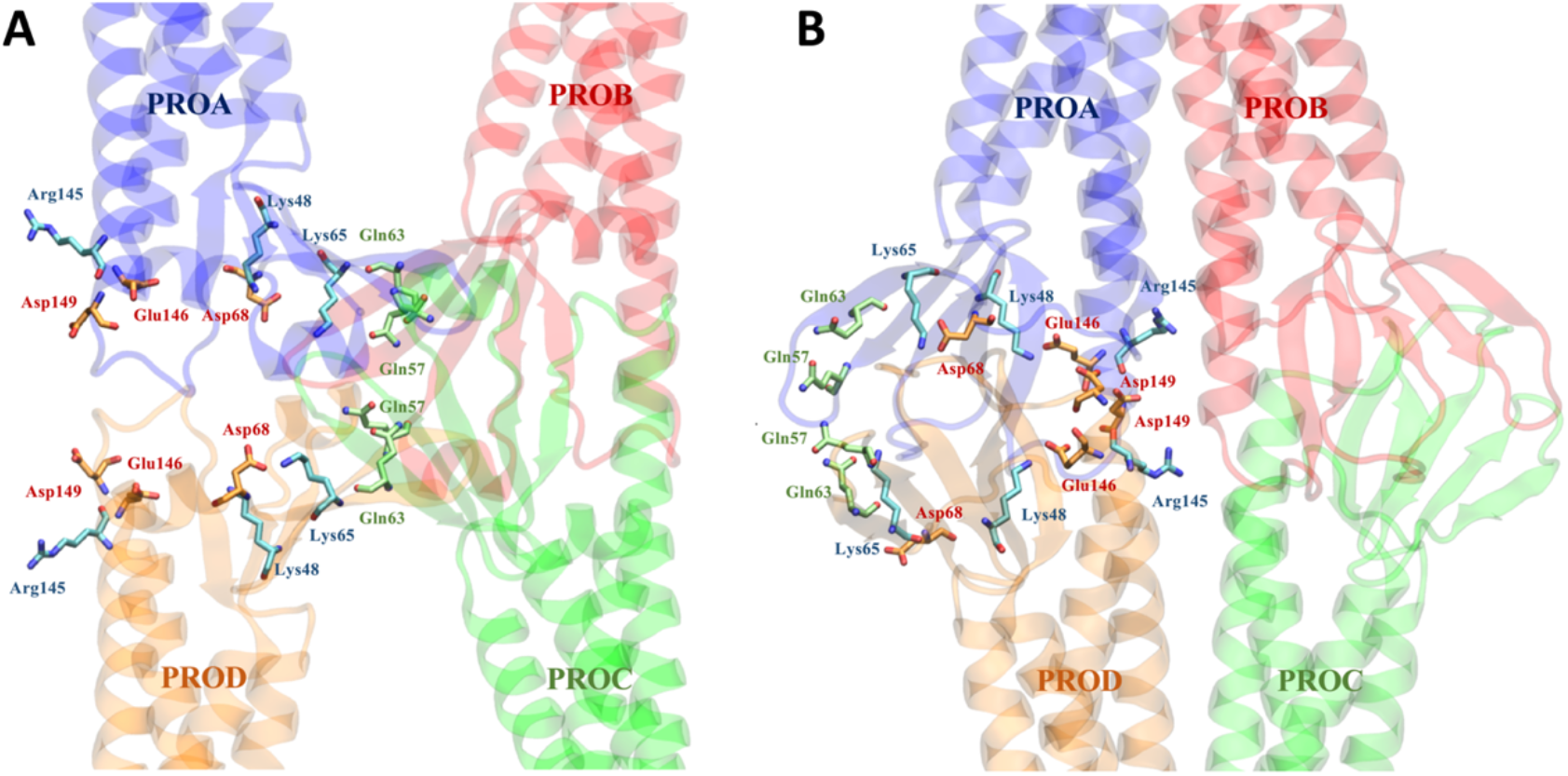
Positions of selected residues along Pore I (A) and Pore II (B) models.

To construct the Pore I system, two distinct models were set-up and simulated. Both the configurations showed a remarkable structural stability of the paracellular domains, evaluated by root-mean square deviation (RMSD) and cross-distances between facing, pore-lining residues. We then selected the model to be used as Pore I system based on the pore size and the preservation of a hydrogen bond involving the highly conserved Lys157 that was described as structurally relevant in Refs. 7,12. Details of the modeling steps and MD simulations set-up are provided in the *Methods* section, while the analysis of the simulations and the assessment of the best Pore I model are reported in the *Supplementary Information* file.

### FE Calculations

Experimental evidence confirms that Cldn5-based TJs form an efficient barrier to the permeation of small molecules and physiological ions^3,6,39,44–47^. Here, to assess the validity of the Cldn5 Pore I and Pore II configurations, we used the Umbrella Sampling (US) method^48^ to perform FE calculations for a single water molecule or single Na^+^, K^+^, Cl^-^, Ca^2+^, Mg^2+^ ions permeating across the cavity of the structures.

In all the US simulations, we used the projection of the position of the tagged ion (or water molecule) on the pore axis as *collective variable* (CV). The FE profiles obtained for the two pore models are reported in **Figure 4** and **Figure 5**, respectively. The errors associated with these calculations were estimated via bootstrapping^49^.

**Figure 4.**
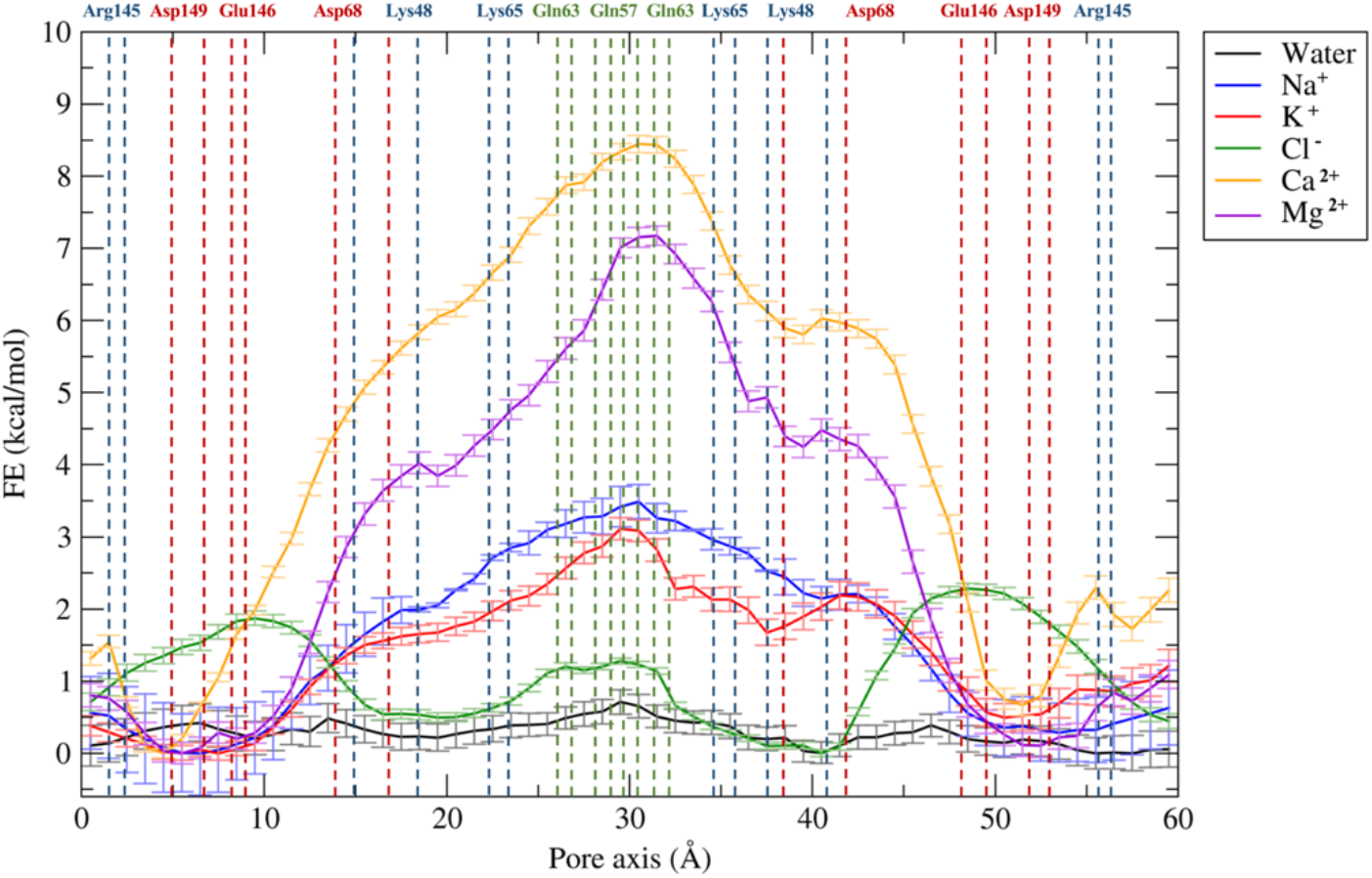
FE profiles for the permeation of water and physiological ions through the Pore I model. The position of the most external atoms of the sidechains of relevant residues are indicated as dashed lines.

**Figure 5.**
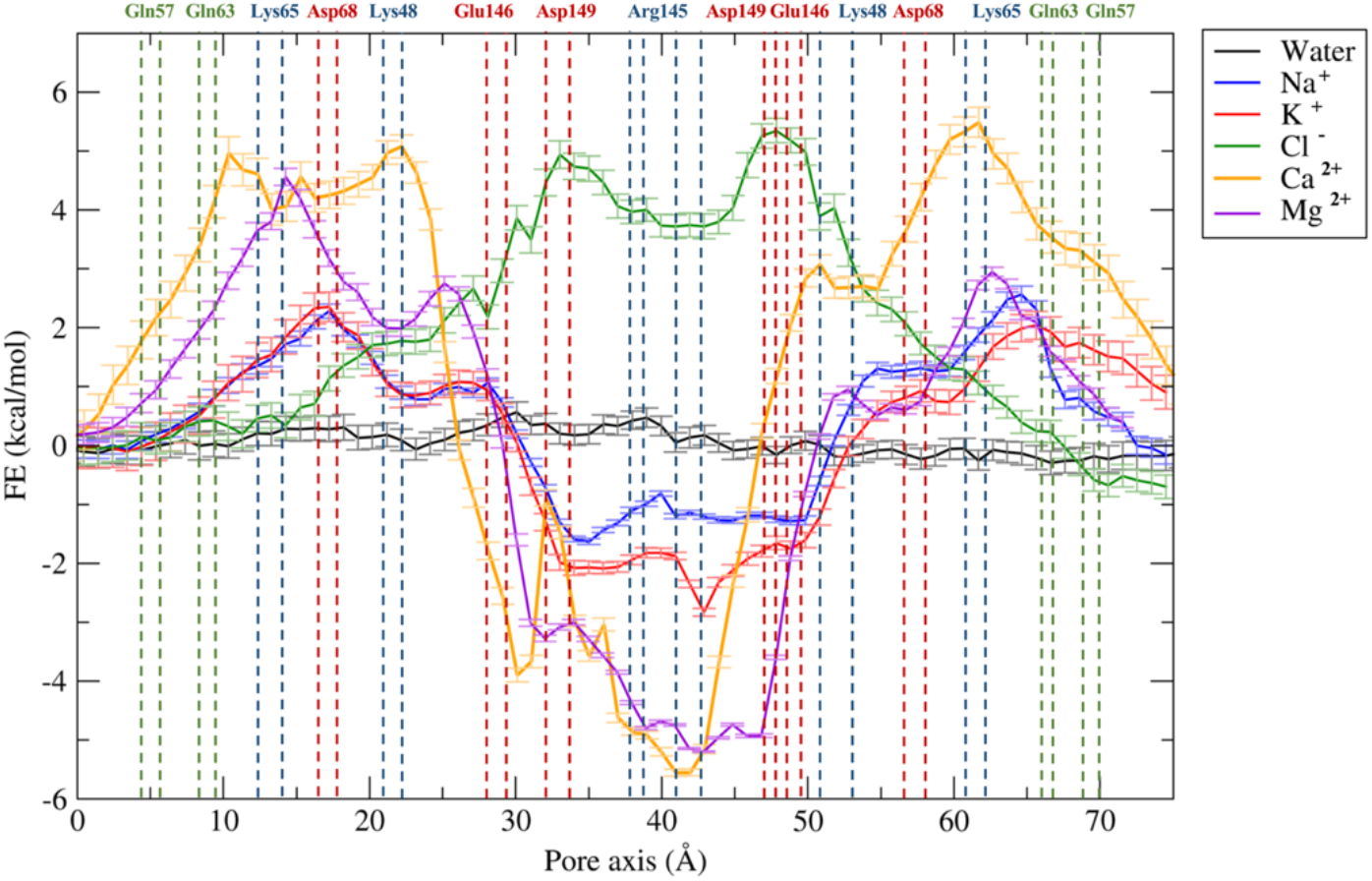
FE profiles for the permeation of water and physiological ions through the Pore II model. The position of the most external atoms of the sidechains of relevant residues are indicated as dashed lines.

The Pore I configuration is characterized by an hourglass shape, with a narrow domain in the middle of the structure, where the Gln57 and Gln63 residues from the four protomers form an uncharged cage (**Figure 3A**). The pore scaffold is completed by positively charged Lys65 residues, which provide an electrostatic barrier to cations. According to the FE profiles illustrated in **Figure 4**, the system is permeable to water. In contrast, the profiles of monovalent (Na^+^ and K^+^) and divalent (Ca^2+^ and Mg^2+^) cations reveal a FE maximum in the constricted region of ^~^ 3 kcal/mol and ^~^ 7-8 kcal/mol, respectively, consistent with the pivotal role of electrostatics in controlling the paracellular transport^33,37,43,50–53^. Overall, these calculations suggest that the Pore I configuration acts as a seal against the paracellular transport of cations. The FE profile for the Cl^-^ ion shows barriers of about 2 kcal/mol symmetrically positioned at the pore entrances. In this regions, two identical clusters of negatively charged residues, Asp68, Glu146 and Asp149 (**Figure 3A**), exert a moderate charge repulsion that limits anion access. Our FE profiles are in overall agreement with those calculated by Irudayanathan et al. ^37^ for the same ions permeating through Cldn5 Pore I (there, the authors used the GROMACS code^54^ and the CHARMM36m force field^55^ with virtual site parameters for lipids^56^, and Well-tempered Metadynamics^57^ for enhanced sampling).

The Pore II system (**Figure 5**) is also water permeable, but it is characterized by a locally different response to ionic transport. The FE profiles for Na^+^ and K^+^ reveal two FE maxima of ^~^ 2 kcal/mol at the two entrances, where the positively charged residues Lys65 and Lys48 are located (**Figure 3B**), together with Asp68. The profiles for Ca^2+^ and Mg^2+^ permeations are characterized by higher barriers, up to 5 kcal/mol. Between the two lateral peaks, a minimum for all cations can be found at the center of the structure correlating with a relevant population of negatively charged residue belonging to the four Cldn5 subunits (Glu146 and Asp149). Because of this cluster of residues, the passage of the Cl^-^ ion is prevented by the presence of a FE barrier reaching 5 kcal/mol that is only slightly damped in the most central region by the four Arg145 residues.

### Pore Size and Hydration of Na^+^ and Cl^-^ during Permeation

To further investigate the link between the FE profiles and the structure of the pores, because ion permeation can be influenced by a combination of steric and electrostatic effects^33^, we calculated the size of the two paracellular cavities. As shown in **Figure 6**, the two models share the same dimension at the two mouths with a diameter of ^~^ 16-18 Å. On the contrary, the internal radius profile differs between the two models. Indeed, the Pore I structure is characterized by an hourglass shape, with an inner constriction in the central part of ^~^ 5-6 Å (**Figure 6A**), where Gln57, Gln63 and Lys65 residues of the four subunits form a narrow cage. On the contrary, the equilibrated Pore II structure has two constrictions of ^~^ 6 Å (**Figure 6B**) in each of the two entrances, where the aforementioned residues belonging to two subunits are located.

**Figure 6.**
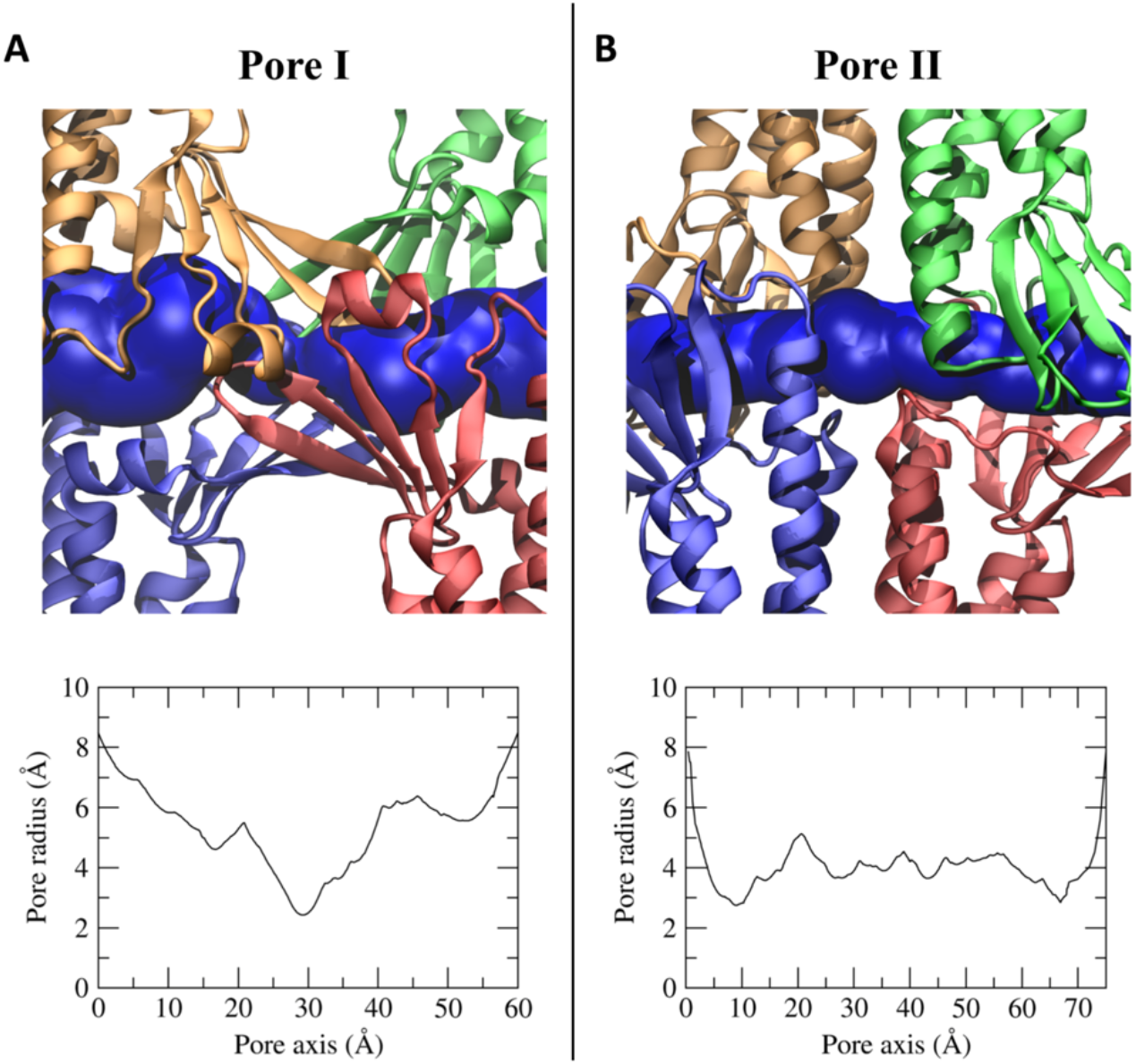
Pore cavities of Pore I (A) and Pore II (B) models. Each protomer is represented in different colors and the pore cavity is shown as a blue surface. In the bottom panels, the pore radius along the pore axis is reported.

We then mapped the hydration pattern of the Na^+^ and Cl^-^ ions during their permeation across the pore cavity (**Figure 7**). To this aim, we calculated the average number of coordinating oxygen atoms belonging to the water molecules surrounding the ions in each US window. For this analysis, we adopted a threshold of 3.0 Å for the cation and 3.5 Å for the anion^58^.

**Figure 7.**
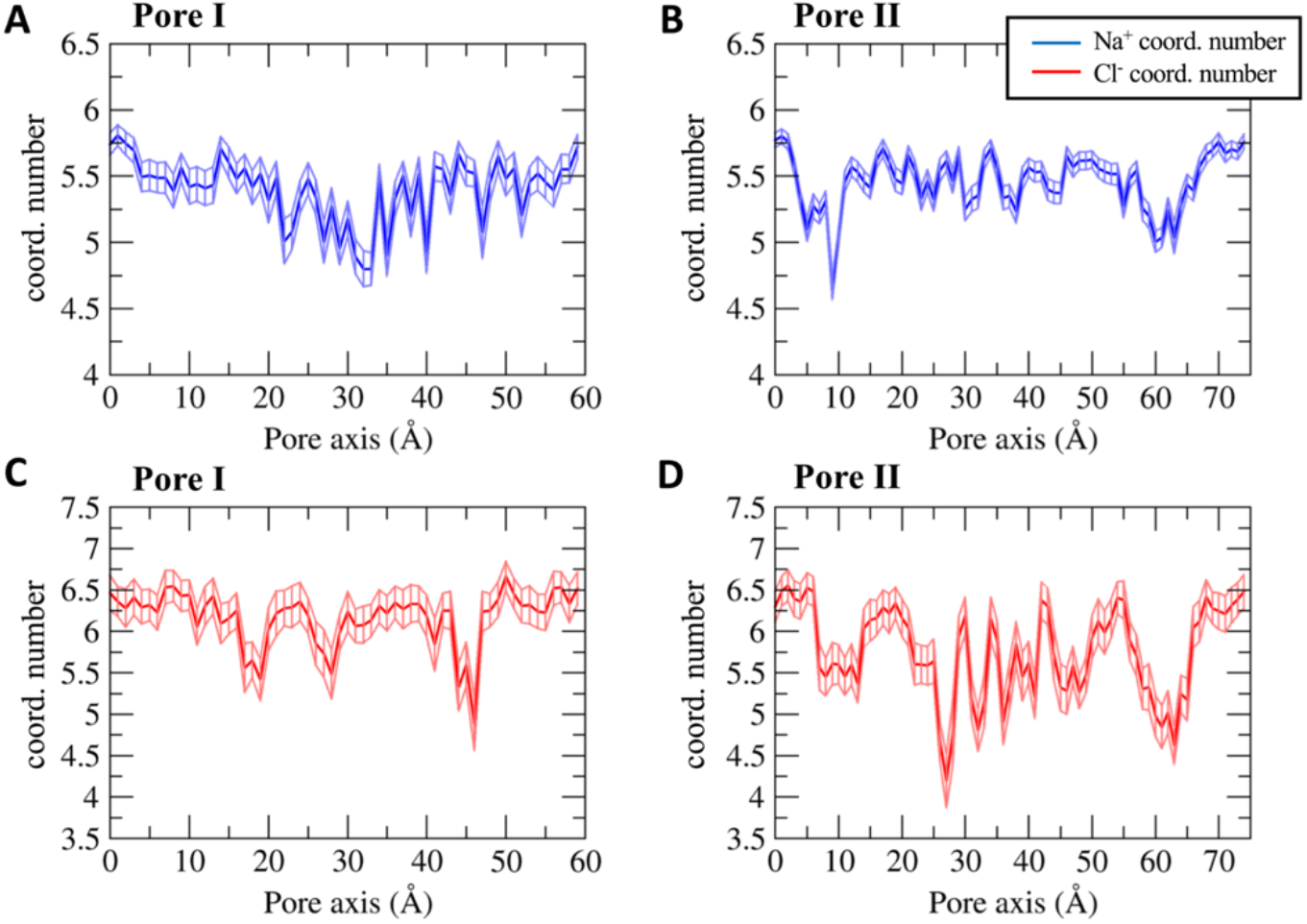
Average number of ion-coordinating water molecules as a function of the pore-axis coordinate for the Na^+^ ion through Pore I (A) and Pore II (B) and for the Cl^-^ ion through Pore I (C) and Pore II (D). Error bars represent the standard deviation of the mean.

The ionic hydration profiles correlate with the pore radius and with the FE profiles of the two structures. The Na^+^ and Cl^-^ ions in the solvent bulk are surrounded by ^~^5.5 and ^~^6.5 water molecules, respectively, in agreement with the values reported in Ref. 58.

The Na^+^ permeating the Pore I cavity (**Figure 7A**) loses up to one coordinating water molecule in the inner region, where the Pore I exhibits the minimal pore radius (**Figure 6A**) and the pore-lining neutral Gln57 and Gln63 residues are located. The partial depletion of the solvation sphere and the unfavorable electrostatic interactions with the positively charged Lys65 residues add up to generate the energetic barrier displayed in **Figure 4**. Similarly, the cation permeating the Pore II cavity (**Figure 7B**) shows two minima in the hydration profile at ^~^10 Å and ^~^60 Å along the pore axis, which form the narrowest regions (**Figure 6B**), where the Lys65 residues are located (**Figure 3B**). This evidence is consistent with the position of the energetic barriers computed with the US calculations (**Figure 5**). Minor fluctuations of the average number of the coordinating molecules are found between the two peaks, where the pore radius (**Figure 6B**) is slightly larger than the radius of the Na^+^ hydration sphere. The hydration profile of the Cl^-^ ion across the Pore I (**Figure 7C**) also correlates with the pore radius and thermodynamics calculations. The FE barriers are found at ^~^10 Å and ^~^50 Å, corresponding to the positions of Asp149 and Glu146. Here, the pore width allows full hydration of the ion, thus partially screening the interaction with the negatively charged residues. Between these regions, three minima at ^~^18 Å, ^~^30 Å and ^~^45 Å are observed in the hydration pattern, correlating with the position of the pore-lining residues Lys48 and with the maximal constriction of the cavity. These findings suggest that the main factor responsible for the formation of the Cl^-^ energetic barriers is the electrostatic repulsion exerted by the negatively charged Asp149 and Glu146 residues, rather than the steric hindrance of the pore. Indeed, in the inner part of the pore, the anion passes through the narrowest segment experiencing a partial dehydration which is not associated to a significant thermodynamic barrier. In contrast, the regions where the FE profiles show the highest barriers to passage of the Cl^-^ are wide enough to accommodate the anion with its entire hydration sphere. The antagonistic contributions of the pore shrinkage and the electrostatics justify the lower entity of the barrier found for the anion (^~^2 kcal/mol) with respect to the monovalent cations (^~^3-3.5 kcal/mol) in the Pore I configuration.

On the other hand, the hydration scheme of the Cl^-^ ion permeating the Pore II model (**Figure 7D**) reports relevant fluctuations because of electrostatic interactions with the pore-lining charged residues and steric hindrance in the tight regions, where the contact with polar amino acids takes place. The minima at ^~^10 Å and ^~^65 Å correlate with the constrictions of the cavity (**Figure 6B**). Nevertheless, the central section reveals limited fluctuations in the pore radius, in the same range of the Cl^-^ hydration sphere. For this reason, the major role to the fluctuations in the coordination pattern of the anion is attributed to the interactions of the ion with the pore-lining residues. To better investigate the mechanisms of the Cl^-^ hydration profiles within Pore II, we analyzed the changes in the coordinating environment of the anion by mapping the interactions of the ion with the pore-lining positively charged residues and the whole protein (**Figure 8**). Results show that, in the regions at ^~^10 Å and ^~^65 Å, the ion interacts not only with Lys65, but also with other protein atoms, due to the constriction of the cavity. In the segments centered at ^~^22 Å and ^~^46 Å, almost all the interactions with the protein are attributed to Lys48, thus revealing a major role of the residue in coordinating the anion to compensate the partial depletion of its solvation sphere. The central segment of the pore axis reveals a fluctuating pattern where the contacts between the anion and the Arg145 residue are predominant. At the sites around ^~^27 Å and ^~^45 Å, corresponding to a pronounced dehydration of the ion, there is substantial interaction with the protein, albeit not with the Arg145 and Lys48 sidechains.

**Figure 8.**
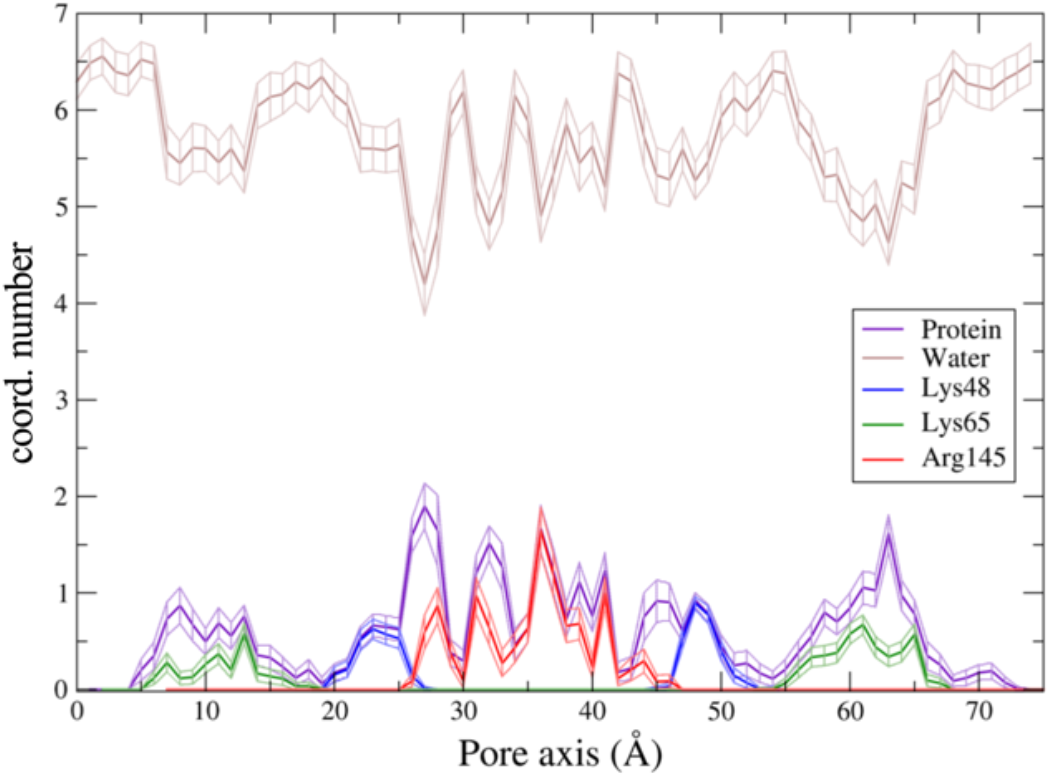
Contributions to the coordination profiles of the Cl^-^ ion in the Pore II cavity. The analysis includes the oxygen atom of the water molecules, the guanidine nitrogen atoms of Arg145, the amine nitrogen atom of both Lys48 and Lys65, the hydroxyl oxygen atom of Ser74 and non-specific heavy atoms of the protein (also including above-mentioned residues).

We next analyzed the time evolution of the hydration pattern in specific US windows, related to representative hotspots in the hydration profile. We mapped the 20-ns-long trajectories of windows 27, 30, 61 and 63 (corresponding to the same positions, expressed in Å, along the pore axis). In window 27 (**Figure 9A**), the Cl^-^ ion loses up to two coordinating water molecules (see also **Figure 7D**). From the analysis of the trajectory, we obtained an average coordination number between the anion and the protein of 1.90 ± 0.24, mainly due to the interaction with the positively charged Arg145 sidechain, the polar Ser74 sidechain and, to a minor extent, the Lys48 residue. Conversely, in window 30 (**Figure 9B**), Cl^-^ is fully hydrated. In this position, we calculated the number of contacts with the negatively charged Glu146 and Asp149 and, as expected, none of them was detected along the entire trajectory. The average coordination number with the protein is only 0.30 ± 0.11, and it is associated with few contacts between the anion and the neighboring positively charged (Arg145) or polar (Ser74) residues. Remarkably, this window corresponds to a high-energy region for the Cl^-^ (**Figure 5**), revealing a major role of the pore-lining negatively charged residues in blocking the anion permeation. Moreover, we investigated the two windows 61 and 63 (**Figure 9 C,D**), where the Cl^-^ ion loses almost two coordinating water molecules. This region of the cavity is one of the most constricted (**Figure 6B**) and it also includes the Lys65. In window 61 (**Figure 9C**), the anion dehydration is mainly due to the stabilizing electrostatic contact with the positively charged Lys65 sidechain, with a coordination number of 0.67 ± 0.10, which represents almost the totality of the anion – protein interaction. In contrast, in window 63, the average coordination number of the protein in contact with Cl^-^ is 1.62 ± 0.19, but the contribution of Lys65 is only 0.34 ± 0.10, revealing that the coordination sphere of the anion is completed by multiple contacts with different polar residues such as Ser58, Gln57 and the Gln63. Consequently, in the segment spanning between 55 Å and 70 Å (and, symmetrically, between 5 Å and 20 Å), the stabilizing electrostatic interaction between the anion and Lys65 cooperates with the steric occlusion and the subsequent contact with the polar sidechains of other pore-lining residues. The most external regions correspond to relatively low-energy values (**Figure 5**), confirming that the electrostatic interactions between the anion and the protein control the permeation process.

**Figure 9.**
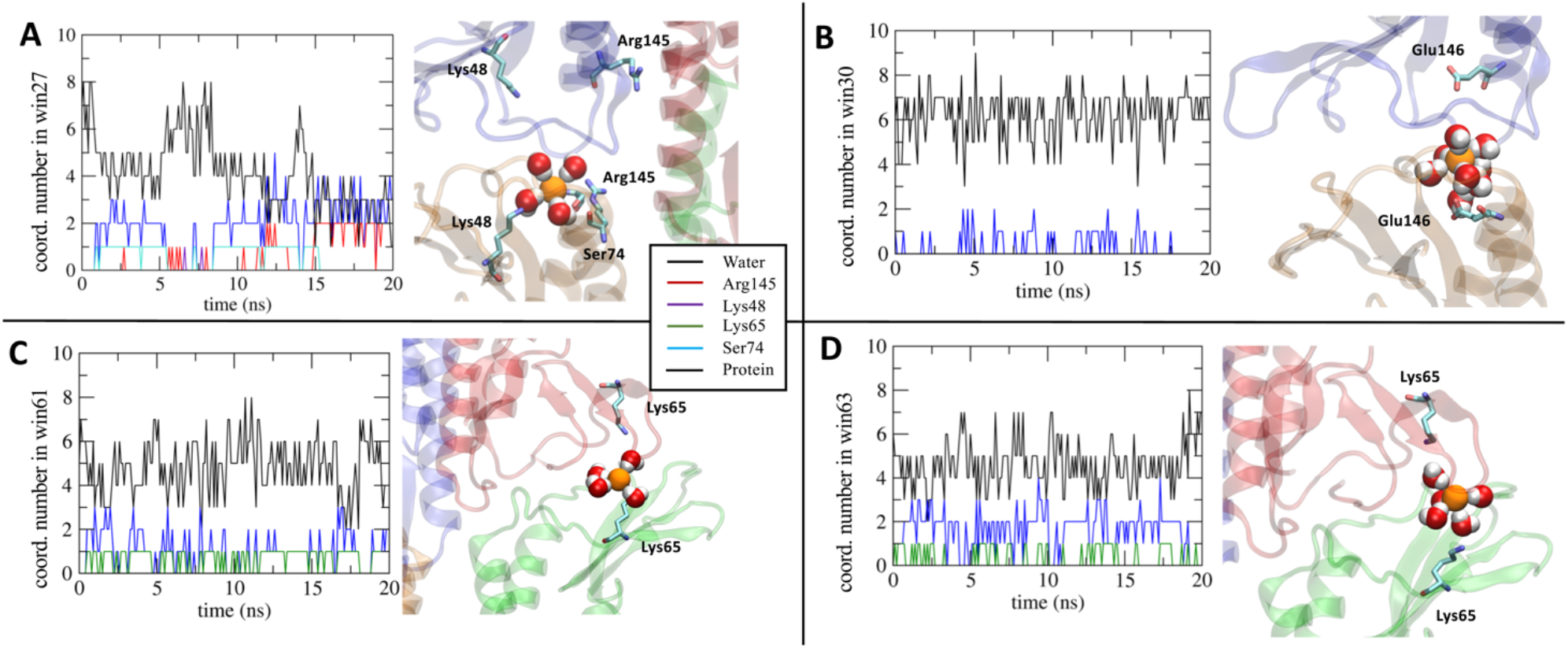
Analysis of the coordination environment of the Cl^-^ ion along the 20-ns-long trajectories in windows (A) 27, (B) 30, (C) 61 and (D) 63 of the US scheme.

### Discussion

Tight junctions are complex intercellular systems observed in both epithelial and endothelial cells, responsible for the control of the paracellular diffusion processes. Among the various tissue-specific TJs, a major interest is devoted to those located in the BBB. Although it is well known that Cldn5 proteins are the backbone of TJ strands of the BBB, we still miss a complete understanding of how they seal the paracellular spaces by oligomerization within the same cell (via *cis* interactions) and between adjacent cells (via *trans* interactions). Here, we refined two structural models of pore-forming Cldn5 complexes that have been recently introduced^40^. These models, proposed for Cldn5 and other Cldns, are only partly consistent with experimental results, so that and their validation has not already been concluded^37,40^. Pore I is in agreement with the structural model proposed by Suzuki and collaborators for the homologous Cldn15^29^, in which two protomers belonging to the same cell interact with their ECL1 domains to form a *cis*-dimer. A couple of these dimers from two opposite cells are then supposed to interact in *trans* forming a Cldn tetramer characterized by a β-barrel-like paracellular cavity^29^. After the publication of this model, criticisms were expressed about its validity^12^, regarding steric hindrances at the paracellular interface and inconsistency with the experimentally demonstrated interactions between TM helices of *cis*-protomers^27,28,42^. On the contrary, in the Pore II structure, Cldn5 *cis*-dimers are formed via a leucine zipper pattern belonging to TM2/TM3. In particular, TM2/TM3-mediated interactions have been previously described by experiments of Cldn5-based systems^28^.Several successive works revised the two models. Various research groups successfully refined the Pore I configuration for Cldn15 and other Cldns^31–38^, showing that non overlapping conformations for the ECLs are possible and that the resulting tetramer is stable. Moreover, experimental results based on electron microscopy techniques^42,59^ indicated an arrangement compatible with the Pore I configuration for Cldn3, Cldn10b and Cldn11. Additional experiments for Cldn3, Cldn10b and Cldn15 showed that the palmitoyl groups are located in proximity of the TM domains^42,60,61^. This last occurrence raises the possibility that palmitoylation could perturb the tight packing of *cis*-interactions at the TM level, thus favoring the Pore I configuration. On the contrary, the multiscale molecular simulations described in Ref. 62 reported that Cldn5 palmitoylation enhances the probability of the dimeric arrangement that characterizes Pore II over the other possible *cis*-configurations occurring at the TM level. Nevertheless, in the absence of a conclusive experimental result able to discriminate between the two models, both the configurations remain worthy of refinement and study, also in the hypothesis of a heterogeneous distribution of the two pore arrangements. Indeed, computational works indicate the potential coexistence of the two pores in the same TJ strands. Coarse grained MD simulations of self-assembly of Cldn protomers suggest diverse possible *cis*-dimers arrangements in the strand, all consistent with the units that construct the two pore configurations^34,35,40^. Furthermore, the same authors confirmed the formation of similar dimers using the PANEL software to obtain millions of Cldn–Cldn conformations and to analyze the amino acid contact maps^41,63^. The results showed the presence of both the Pore I and Pore II structures investigated in this work for Cldn5.

In this framework, we used MD simulations and FE calculations to quantify the thermodynamic features of ionic permeation events through the pore cavities of the two Cldn5 models. The study of ionic processes across biological channels has been an important topic in molecular modeling to look into the details of protein models^64^. In our previous work, we used the same approach to refine the configuration of the Cldn15 pore based on the original structure of Ref. 29. Our efforts contributed to the validation of this structure, confirming the role of the investigation of ion permeation processes for structural validations. In this work, we extended our analysis to the two structural models built for Cldn5 subunits. The HOLE profiles revealed a different pore shape between the two models. The Pore I structure is characterized by an hourglass shape, with the inner constriction in correspondence of residues Gln57, Gln63 and Lys65 of the four subunits, measuring ^~^ 5-6 Å in diameter. On the contrary, the Pore II structure has two constrictions of ^~^ 6 Å in diameter each in proximity of one of the two entrances, where the same residues Gln57, Gln63 and Lys65, now from two subunits, are located. Indeed, it is remarkable that, despite their different topologies, the narrowest pore regions in both models are related to the same set of residues. Our FE calculations reveal that both the pores are water permeable, a feature not yet fully clarified experimentally, but consistent with previous computational results^37,40^ and postulated by some authors^65^. On the contrary, the pores show FE barriers to cations. Interestingly, in both the models, the position of the cation barriers corresponds to the narrowest regions. This is in line with the fact that the minimum pore diameters are close to the size of a hydrated Na^+^ ion, and slightly smaller than the diameter of the Cl^-^ hydration shell. This observation pushed us to investigate the details of ion hydration during Na^+^ and Cl^-^ permeation by US simulations. The passage of the Na^+^ ion through the constrictions induces a partial dehydration of its shell. This contributes to generate the FE barrier, together with the unfavorable electrostatic repulsions between the cation and the Lys65 residues of the different Cldn5 subunits. The coupling of steric and electrostatic effects has been already observed in another Cldn-based paracellular system^33^ and it is also relevant for the study of other, more conventional, narrow protein channels such as gramicidin^66^ or in the selectivity filter of K^+67,68^ and Na^+69^ channels. As for the hydration pattern of Cl^-^, both steric and electrostatic effects induced pronounced fluctuations in the number of water molecules coordinating the anion passing through the pore axis. The smaller solvation energy of Cl^-^ with respect to Na^+^ (−6.4 kcal/mol and −17.2 kcal/mol, respectively^70,71^) provide a more labile coordinating shell to the anion, and the effects are particularly evident in the regions of the cavities where the pore radius is comparable to the size of the Cl^-^ hydration sphere. The analysis of the coordinating environment of the anion permeating the Pore II cavity revealed that the energetics of the barriers are mainly driven by the unfavorable electrostatic interactions with the pore-lining acidic residues, while the depletion of the solvation sphere due to steric hindrance is not correlated to high energy regions. Conversely, a stabilizing interacting network with positively charged and polar amino acids able to fill the solvation sphere of the ion is observed in the regions of maximal constriction. For both the models, the FE profiles suggest that the permeation of Cl^-^ is limited by the presence of the negatively charged Asp149 and Glu146 residues. These data indicate the absence of preference for cation *versus* anion selectivity for both the pore models, in agreement with the well-known characteristics of the BBB^20,44,47^. Our results complement those exposed for the Pore I in Ref. 37, extending the validation of the two models in terms of ionic permeation features.

### Conclusions

The BBB plays a pivotal role in controlling the brain homeostasis, thanks to its high selectivity that prevents the passage of harmful molecules from the blood. As a consequence, it is a significant obstacle to effective brain drug delivery in the treatment of CNS diseases^15–17,20,72–74^. To overcome this limitation, strategies are emerging to enhance the BBB permeability by modulating the passive transport across the TJs in the paracellular space^21–25^. This approach has already provided promising results from *in-vitro* experiments of drug-enhancer peptides^75,76^. Although it is well-known that the TJ scaffold is essentially formed by Cldn5 protein complexes, the fine structural details of Cldn5 multimeric arrangement are still missing^7,12,77^.

Recently, two tetrameric pore-forming models have been introduced after computational investigations based on coarse grained MD simulations^40^. Despite the different topological configurations, both the structures, originally named Pore I and Pore II, recapitulate various features from experimental results^29,30,78^, but a systematic comparison of these systems is still missing. In this work, we refined the structures of the two Cldn5 pore configurations in solvated double-bilayer environment by all-atom MD simulation. Then, we calculated the FE profiles for single-ion translocation across the two pores. Both the structures fit the typical barrier-like behavior of Cldn5 in the BBB TJs. The findings illustrated in this work extend our knowledge of Cldn5 TJ structures and, although in the simplest case of single-pore systems, offer a molecular description of the BBB Cldn5 role. Furthermore, by identifying Cldn5 homomeric interaction surfaces in the TJs, our results can contribute to develop experimental strategies to enhance the drug delivery process across the BBB by modulating the paracellular permeability.

## METHODS

### Pore I

The Pore I configuration was assembled with four Cldn5 protomers matching the quaternary structure published by Suzuki et al.^29^. The Cldn5 protomers were modeled from the Cldn15 homologs using structures from two different works^30,32^, obtaining two putative models for the Pore I. The first model (named *Model1*) was built starting from the tetrameric configuration of the Cldn15 simulated by Alberini et al.^32^ The second one (named *Model2*) was assembled starting from the configuration of the Cldn15 pore published by Suzuki et al.^29^. In this case, a single Cldn5 structure was modelled adopting the crystal structure of the mouse Cldn15 protomer as a template (PDB ID: 4P79)^30^.

#### Model1

The pore simulated by Alberini et al.^32^ was disassembled in four separated Cldn15 monomers which have adopted slightly different conformations after the simulated trajectories of 250 ns described in Ref. 32. Each of these four protomers were used for the homology modelling of four Cldn5 monomers via the SWISS-MODEL^79^ program. The four raw models of Cldn5 were then refined with the ModRefiner^80^ server. The resulting protomers of Cldn5 were superimposed on the Cldn15 template^32^ with the UCSF Chimera^81^ Matchmaker tool. Afterwards, the tetrameric system was refined with the GalaxyRefineComplex tool^82,83^ and the configuration with highest score was selected for MD simulations. The complex was then oriented with the pore axis parallel to the cartesian y-axis and embedded in a double bilayer of pure 1-palmitoyl-2-oleoyl-sn-glycero-3-phosphocholine (POPC), solvated with explicit three-point (TIP3P)^84^ water molecules and charge-neutralized with counterions using VMD 1.9.3^85^. The fully-hydrogenated pdb file of the protein complex was generated with the CHARMM-GUI PDB manipulator tool^86,87^. Two hexagonal membranes were generated using the *membrane builder* tool of the same platform^87,88^ and equilibrated separately for 10 ns with the NAMD 3.0 software^89^ and the CHARMM36m force field^55^ using hexagonal periodic boundary conditions. The final simulation box is a hexagonal prism with a base inscribed in a square of approximately 120.0 × 120.0 Å^2^ and a height of around 160.0 Å. The topology file was built with the psfgen tool of VMD 1. 9.3^85^ with the parameters of the CHARMM36m force field^55^ and the four disulfide bridges were preserved between residues Cys54 and Cys64 found in the ECL1 of each protomer.

#### Model2

The Cldn5 protomer for Model2 was built starting from the crystal structure of the isolated Cldn15 published by Suzuki et al. (PDB ID: 4P79)^30^. The crystal lacks a segment of eight residues (34–41) in the ECL1 that is automatically built by SWISS-MODEL^79^ during the homology modelling of Cldn5. The resulting structure was refined with ModRefiner^80^, consistently with the workflow illustrated for the Model1 and replicated in four identical copies. Following the same protocol illustrated for Model1, the four Cldn5 protomers were assembled to form the tetrameric arrangement. Analogously, the optimal system was embedded in a hexagonal double POPC bilayer, solvated with water and charge-neutralized with counterions.

#### Equilibration and unbiased MD simulation

Both Model1 and Model2 systems contain about 200000 atoms. They were equilibrated with a multi-step protocol where, after a first energy minimization, they were heated up to 310 K and simulated for 30 ns with a progressive release of positional restraints on the heavy atoms. Each model was then simulated for 1μs. To avoid any rigid body rotational or translational displacement of the protein, the coordinates of the C*α* atoms of the residues 6, 9, 20, 23, 79, 82, 97, 100, 117, 120, 138, 141, 166, 169, 177, 180, all belonging to the most external residues on the TM *α* helices, were restrained to their initial values by harmonic potentials. The use of restraints on TM and/or ECLs backbone atoms in the isolated pore conformations can be justified by the fact that this model structure misses the neighbor protomers of the strands, which in the physiological TJ architecture form a scaffold that constraints the pore, limiting the fluctuations of its domains. Notably, in all of our extended MD simulations of Pore I the ECL domains were stably preserving the *β*-barrel structure, and so we limited the restraints to few atoms of TM helices. The systems were simulated in the NPT ensemble at P= 1 atm and T= 310 K, maintained by a Langevin thermostat and Nosé-Hoover Langevin piston^90,91^. Long range electrostatic interactions were computed using the Particle Mesh Ewald (PME) algorithm^92^. Chemical bonds between hydrogen atoms and protein heavy atoms were constrained with SHAKE^93^ while those of the water molecules were kept fix with SETTLE^94^. The NAMD 3.0 program^89^ with CHARMM36m force field^55^ was used to perform the simulations. Based on measured pore size and the presence of a hydrogen bond deemed structurally relevant in Refs. 7,12, we selected Model2 to represent Pore I and continue with the FE calculations. The comparative analysis of the simulations are reported in the *Supplementary Information* file.

### Pore II

The Pore II configuration was described in Ref. 40. In this architecture, the *cis*-interface originates from to the interaction of two neighboring protomers at the level of the TM helices arranging in a leucine zipper composed by the residues Leu83, Leu90, Leu124 and Leu131 on the TM2 and TM3, supported by two homophilic *π-π*-interactions between Phe127 and Trp138 on the opposing TM domains (**Figure 1B**). To build this structure, we first simulated a Cldn5 protomer, again homology-modeled from the Cldn15 template. The protein was embedded in a rectangular pure POPC membrane bilayer and equilibrated with a 110-ns-long all-atom MD simulation in explicit solvent. The trajectory was analyzed to assess the structural stability of the protein (see *Supplementary Information* file). An equilibrated configuration of the Cldn5 protomer was extracted from the trajectory and used to reproduce the *cis*-interface first via a docking protocol. The leucine zipper TM interaction between two copies of the protein was predicted by the MEMDOCK server^95^, which includes a specific algorithm for docking *α* – helical membrane proteins. The dimer selected by MEMDOCK^95^ was further refined with DOCKING2^96–98^, and the structure finely reproducing the leucine zipper was embedded in a pure POPC membrane, solvated with explicit water and equilibrated with ^~^100 ns of all-atom MD simulation in presence of charge-neutralizing counterions. The final dimer complex was replicated and the two copies were used to assemble the Pore II configuration with a further docking approach. Following the protocol suggested in Ref. 40, the Pore II complex was generated using ClusPro^99–103^ to reproduce the *trans*-interactions occurring between two opposing dimers at the level of the paracellular domains. Afterwards, the tetrameric structure was refined using GalaxyRefine^82,83^, oriented with the pore axis parallel to the cartesian y-axis and embedded in a hexagonal double bilayer of pure POPC, solvated with water and charge-neutralized with counterions. The topology file was built with the psfgen tool of VMD 1.9.3^85^ with CHARMM36m parameters^55^ and the four disulfide bridges were preserved between residues Cys54 and Cys64 found in the ECL1 of each protomer.

#### Equilibration and unbiased MD simulation

The Pore II simulation set-up (^~^200000 atoms) followed the same protocol described for the two putative models of Pore I. Additionally, further harmonic restraints were applied on the C*α* atoms of the residues 11, 14, 25, 28, 78, 81, 99, 102, 116, 119, 143, 146, 166, 169, 183, 186 in the ECLs.

### Pore size analysis

The size of the paracellular pores was monitored along the trajectory with the HOLE program^104,105^. The algorithm maps the radius of a protein cavity along a given axis (here, the y-axis) by fitting a spherical probe with the Van der Waals radii of the pore-lining atoms. For all the models, a 15 Å threshold was chosen for the pore radius and representative structures spaced by 10 ns along the production trajectory were selected and analyzed (see *Supplementary Information* file).

### Free energy calculations

The free energy (FE) profiles for the permeation of a water molecule and single ions were calculated using the Umbrella Sampling (US) method^48^. A restraining term is added to the MD potential to confine a *collective variable* (CV, function of the Cartesian coordinates of the system) in selected regions, named *windows*, allowing proper sampling even in high-energy regions of the landscape. As CV, we chose the coordinate of the tagged permeating ion along the pore axis, previously aligned with the cartesian y-axis, and the restraining potential *V_i_*(*y*) in each window *i* is:

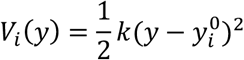

where 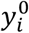 indicates the value in Å at which the CV is restrained in the window (called *center*) and *k* is a constant that is appropriately chosen in order to ensure a sufficient overlap of the CV distributions from adjacent windows (in this work, we used *k* = 2.0 *kcal*/(*mol* Å^2^) for all the simulations). In each window, the displacement of the ion orthogonal to the pore axis is confined within a disk of radius *r*_0_ + *δ*, where *r*_0_ is the pore radius as determined by the HOLE program^104,105^ and *δ* = 2 Å. The equilibrated conformation of the system was used as starting structure of all US windows, and the ion was manually positioned at each center 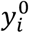. The Pore I cavity was split into 60 windows. Each window was minimized and simulated for 16 ns with the same set up described for the standard MD simulation, adding the bias to the force field^106^ via the *colvors* module^107^. Positional restraints on selected C*α* atoms were applied as described for the unbiased simulations. The first ns of production was excluded from the statistics. Due to the elongated shape of the Pore II cavity, 75 windows were required to sample the entire cavity. The simulation followed the same protocol adopted for the Pore I, except that 20 ns of production per window were carried out in order to achieve a proper convergence. The FE profiles are obtained by combining the CV distributions of all windows using the *weighted histogram analysis method* (WHAM)^49,108,109^. We employed the code from the Grossfield group available at http://membrane.urmc.rochester.edu/content/wham. The *block error analysis* was also implemented to the calculated FE profiles (see *Supplementary Information*).

### Safety

No hazards or risks were associated with this works.

## List of Abbreviations

BBB: Blood-brain barrier
CLDN: Claudin
TM: Trans-membrane
ECL: Extracellular loop
FE: Free energy
MD: Molecular dynamics
RMSD: Root-mean square deviation
US: Umbrella sampling
WHAM: Weighted histogram analysis method

## Author Information

Alessandro Berselli

Giulio Alberini

Fabio Benfenati

Luca Maragliano

*Corresponding Authors:* Fabio Benfenati and Luca Maragliano

## Author Contributions

Alessandro Berselli and Giulio Alberini equally contributed to perform the simulations, analyze the data, discuss the results and write the manuscript. Luca Maragliano and Fabio Benfenati supervised the research project, discussed the results and revised the manuscript.

## Supporting Information

All-atom MD simulation results and analysis of the Pore I models, free energy profiles with the block error analysis and all-atom MD simulation results and analysis of the Cldn5 protomer.

## Acknowledgment

We thank Jörg Piontek and Matteo Ceccarelli for useful discussions; Sergio Decherchi, Diego Moruzzo and Andrea L. Benfenati for valuable help. Computing time allocations were granted by the CINECA supercomputing center under the ISCRA initiative. We also gratefully acknowledge the HPC infrastructure and the Support Team at Fondazione Istituto Italiano di Tecnologia.

---

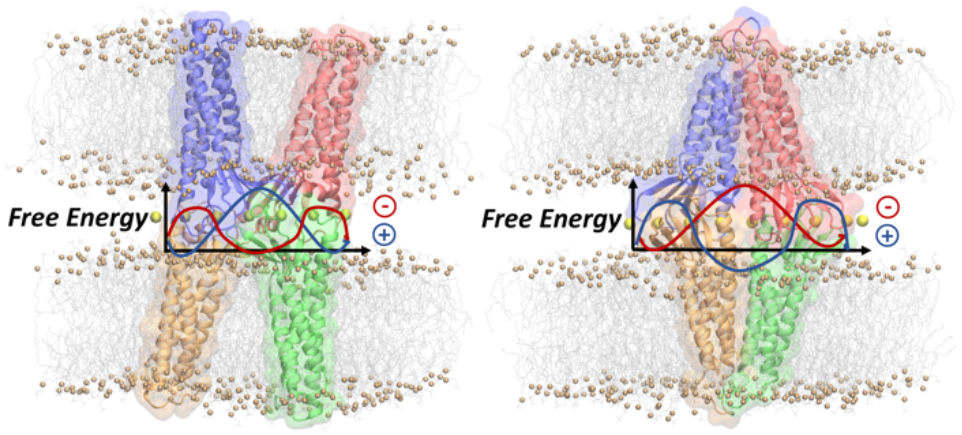
For Table of Contents only. 8.22 cm X 3.92 cm

