## Supplementary Information file for "Computational Assessment of Different Structural Models for Claudin-5 Complexes in Blood-Brain-Barrier Tight-Junctions"

**List of Abbreviations**

- BBB: Blood-brain barrier
- CLDN: Claudin
- TM: Trans-membrane
- ECL: Extracellular loop
- FE: Free energy
- MD: Molecular dynamics
- RMSD: Root-mean square deviation
- US: Umbrella sampling
- WHAM: Weighted histogram analysis method

**List of Models**

**Pore I - Model 1**

Single pore configuration, where the four Cldn5 promoters were homology-modeled based on the equilibrated configuration of the Cldn15 single pore model illustrated in Ref. 1.

**Pore I - Model 2**

Single pore configuration, where each Cldn5 promoter is modeled based on the homology with the crystal structure of the mouse Cldn15 promoter (PDB ID: 4P79)[2]. The configuration of the single pore model is obtained starting from the atomic coordinates of the pdb file provided in Ref. 3.

### CONTENTS

1. ANALYSIS OF STANDARD MOLECULAR DYNAMICS SIMULATIONS OF TWO VERSIONS OF THE PORE I MODEL **S2**

**1.1 RMSD** **S2**

**1.2 Cross-distances** **S2**

**1.3 Pore Radius** **S7**

2. BLOCK ERROR ANALYSIS **S8**

3. STRUCTURAL ANALYSIS OF THE CLDN5 MONOMER **S10**

REFERENCES **S12**

### 1. Analysis of Standard Molecular Dynamics Simulations of two Versions of the Pore I model

#### 1.1 RMSD

We calculated the root mean squared deviation (RMSD) of the atomic backbone with respect to the starting conformation, using configurations sampled every 100 ps along the trajectories. The RMSD was computed for different domains of the structures (**Figure S1**). In the following, we indicate the four protomers using the PROA/B/C/D labels. For both the models, we observe a plateau between 2.5-3 Å for the tetramer, as illustrated in **Figure S1A**. The RMSD for each protomer and for each ECL was calculated (**Figure S1B, S2**). These results show that the integrity of the $\beta$-barrel is preserved, and the dimension of the cavity does not reveal significant variations, as confirmed by the pore radius calculations reported in **Figure S6**.

#### 1.2 Cross-distances

We calculated various cross-distances between pore-lining residues as illustrated in **Figure S3**. In particular, for each pair of residues, we computed the distance between the two C$\alpha$ atoms and between the last C-atom of the sidechains, as summarized in **Table S2**. The Gln57 and Gln63 residues are located in the narrowest region, while Lys65 is next to them (**Figure S3C**). The distance between the residues Val70 and Pro153 accounts for the integrity of the lateral scaffold of the pore while the other between the residues Lys48 describes the fluctuations of the pore diameter at its entrance (**Figure S3D**).

Both the models exhibit stationary fluctuations for each cross-distance, as illustrated in **Figure S4**. In particular, for the Gln63 pairs in Model1 the average value of the CD – CD distance (between the C-atom at the extremities of the sidechains) is 12.29 $\pm$ 1.15 Å for the PROA-PROD dimer, and 8.32 $\pm$ 1.19 Å for the PROB-PROC dimer (**Figure S4A**). These distances are smaller than those found in the Model2, where values of 15.89 $\pm$ 1.28 Å and 13.69 $\pm$ 1.02 Å, respectively, are obtained (**Figure S4B**).

The distances between residues at the mouth of the pore are reported in **Figure S4**.

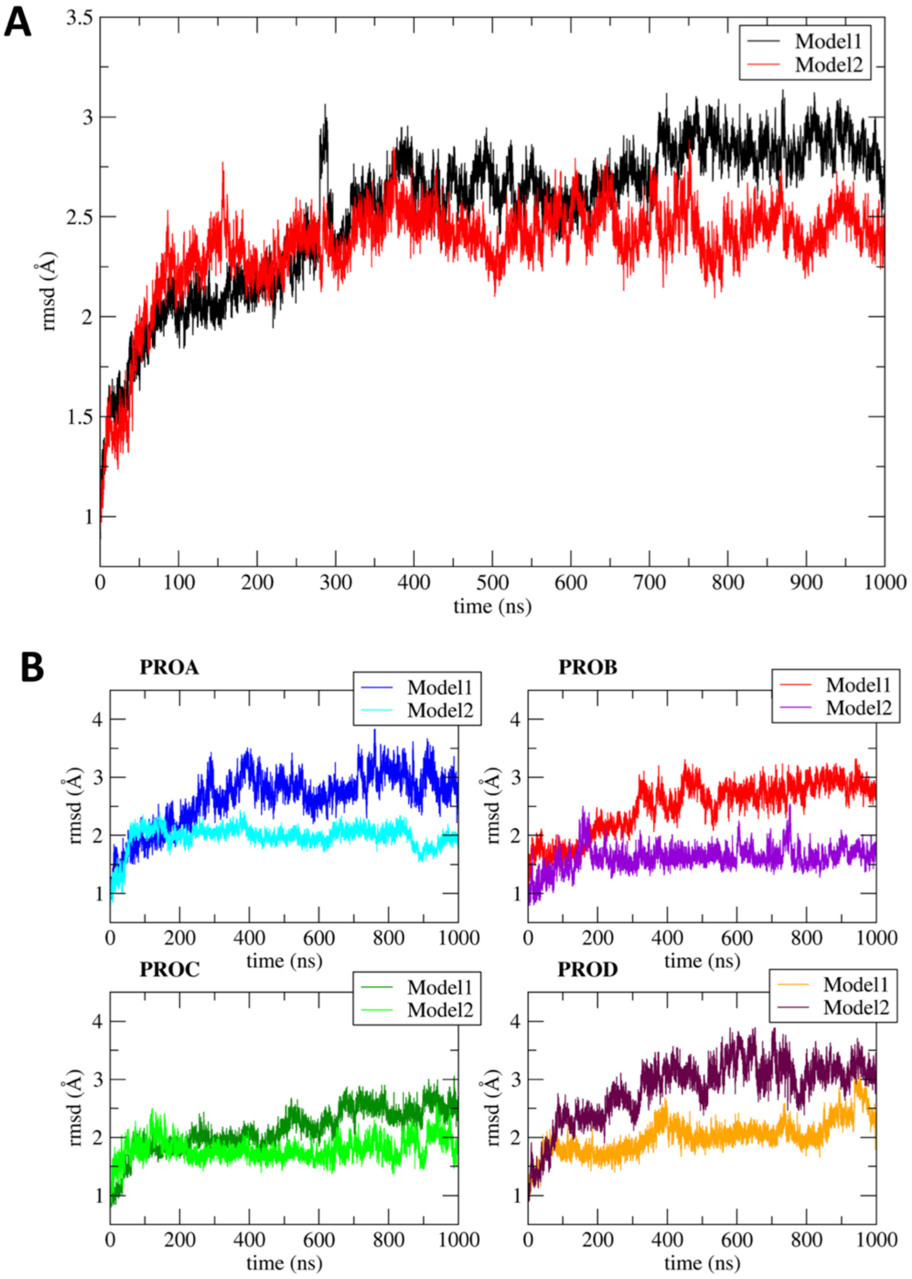


***Figure S1**. Backbone RMSD of Model1 and Model2. A) RMSD for the tetramer. B) RMSD for each protomer.*

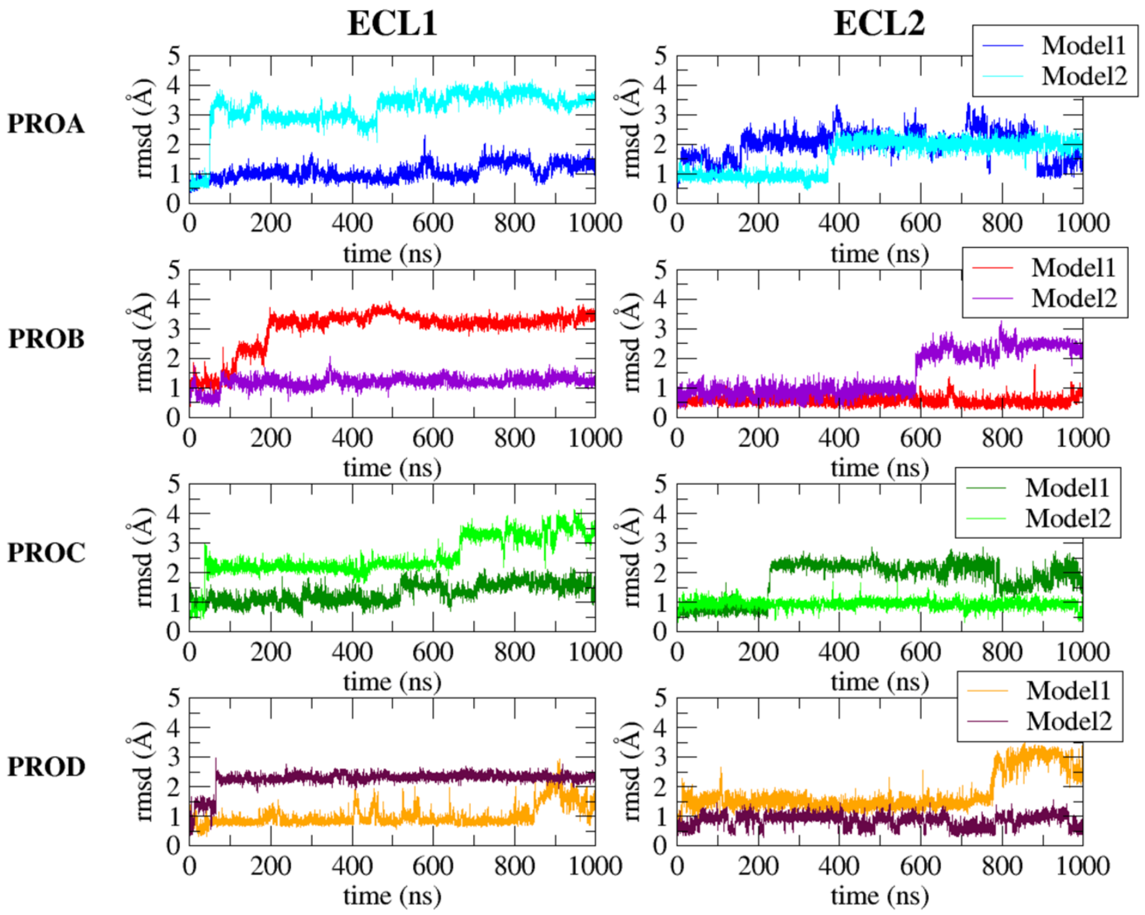


***Figure S2****. RMSD of the Model1 and the Model2 ECLs.*

***Table S1****. Cross-distances mapped along the 1-µs-long MD simulation of the two Pore I models.*

| Residue | Atom type | Protomers |
|---|---|---|
| **Gln57 – Gln57** | CD – CD | PROA - PROD |
| | CA – CA | |
| | CD – CD | PROB - PROC |
| | CA – CA | |
| **Gln63 – Gln63** | CD – CD | PROA - PROD |
| | CA – CA | |
| | CD – CD | PROB - PROC |
| | CA – CA | |
| **Lys65 – Lys65** | CE – CE | PROA - PROD |
| | CA – CA | |
| | CE – CE | PROB - PROC |
| | CA – CA | |
| **Lys48 – Lys48** | CE – CE | PROA - PROD |
| | CA – CA | |
| | CE – CE | PROB - PROC |
| | CA – CA | |
| **Pro153 – Val70** | CG – CB | PROA - PROD |
| | CA – CA | |
| | CG – CB | PROB - PROC |
| | CA – CA | |

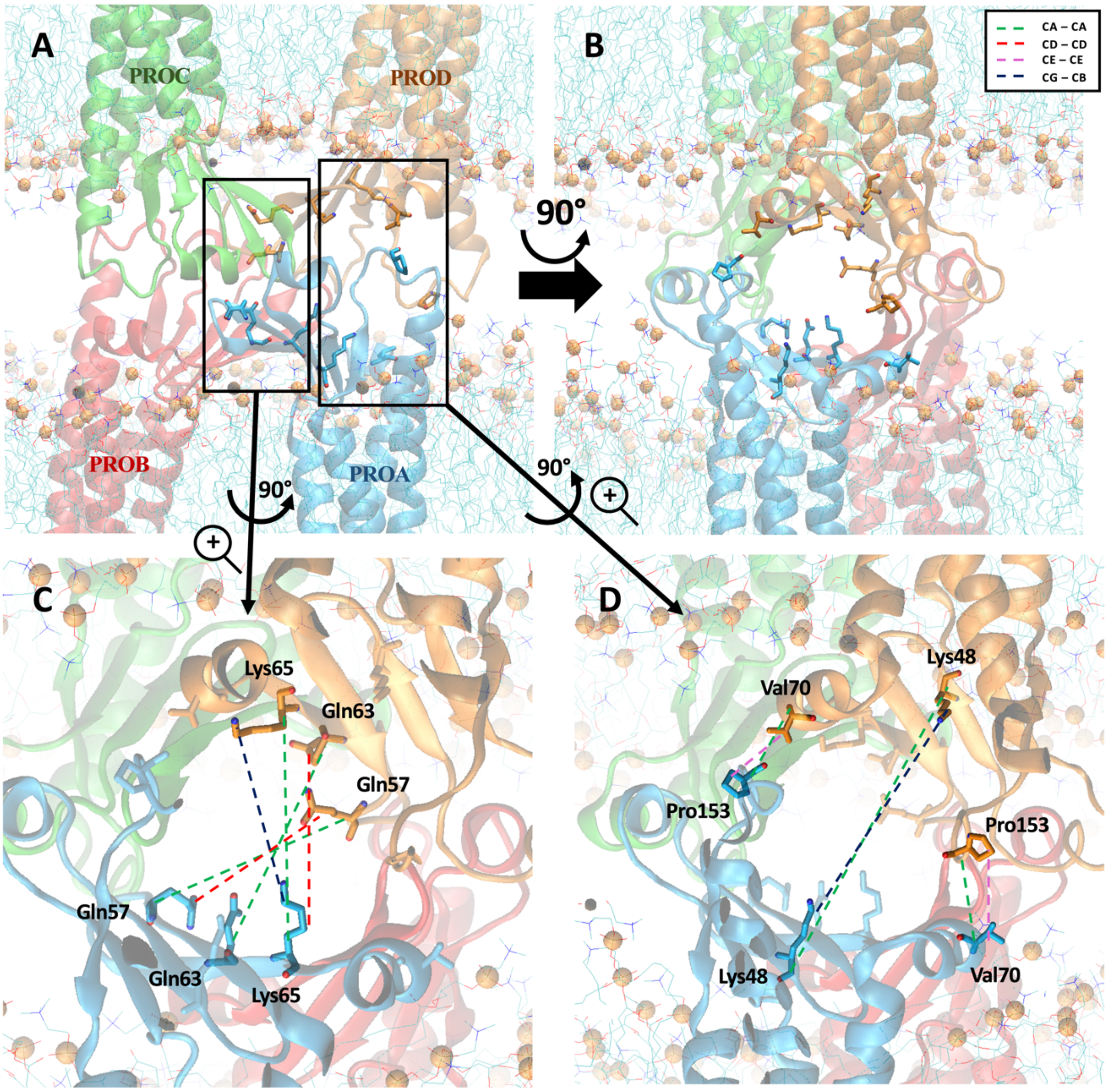


***Figure S3***. *Graphical representation of the cross-distances calculated along the trajectories of Model1 and Model2, visualized on Model2. Individual protomers are distinguished by their coloring and referred as PROA (blue), PROB (red), PROC (green), PROD (orange). C-atoms are reported with the CHARMM atom name. A) Lateral side view; residues are displayed only for protomers PROA and PROD. B) Apical view of the pore cavity. C) Close-up view of the distances monitored at the center of the pore. D) Close-up view of the distances monitored at the mouth of the pore.*

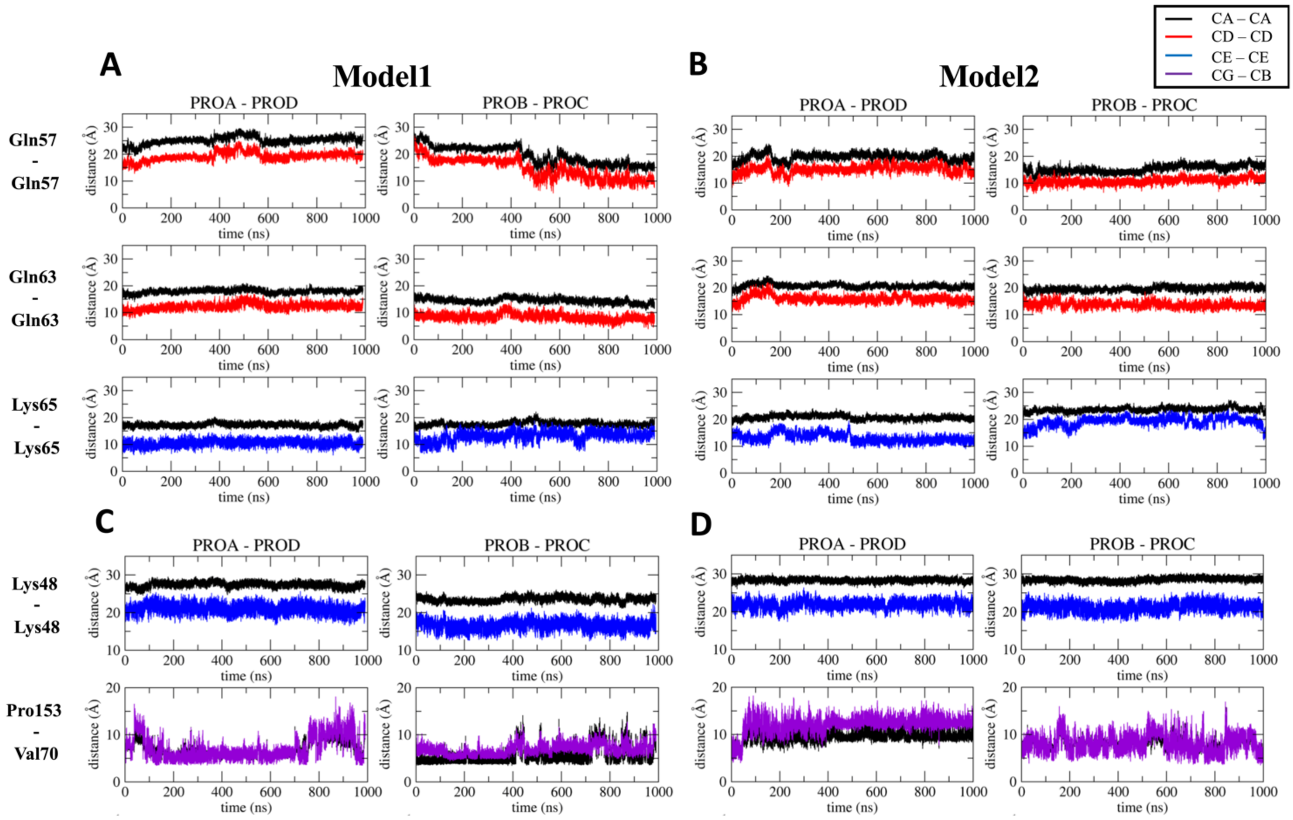


***Figure S4.*** *Cross-distances for Model1 and Model2. A) Residue pairs close to the pore center of Model1. B) Residue pairs close to the pore center of Model2. C) Residue pairs at the entrance of the Model1 pore. D) Residue pairs at the entrance of the Model2 pore.*

In Ref. 4, the authors reported the presence of the hydrogen bond (HB) between the sidechain of Lys155 and the backbone of Asn148 as pivotal for the preservation of the ECL2 rigidity in the structure of the mouse Cldn15 (PDB ID: 4P79)[2]. Consistently, in the present work, we identified the corresponding interaction in the Cldn5-based models between the residues Lys157 and Asp149 (**Figure S5A**). In each model, we monitored the value of the distance between the N-atom of the Lys157 sidechain and the O-atom of the Asp149 backbone. The interaction is not preserved in any of the four protomers belonging to Model1. On the other hand, in Model2, the distances correlate with a HB for three (**Figure S5B**).

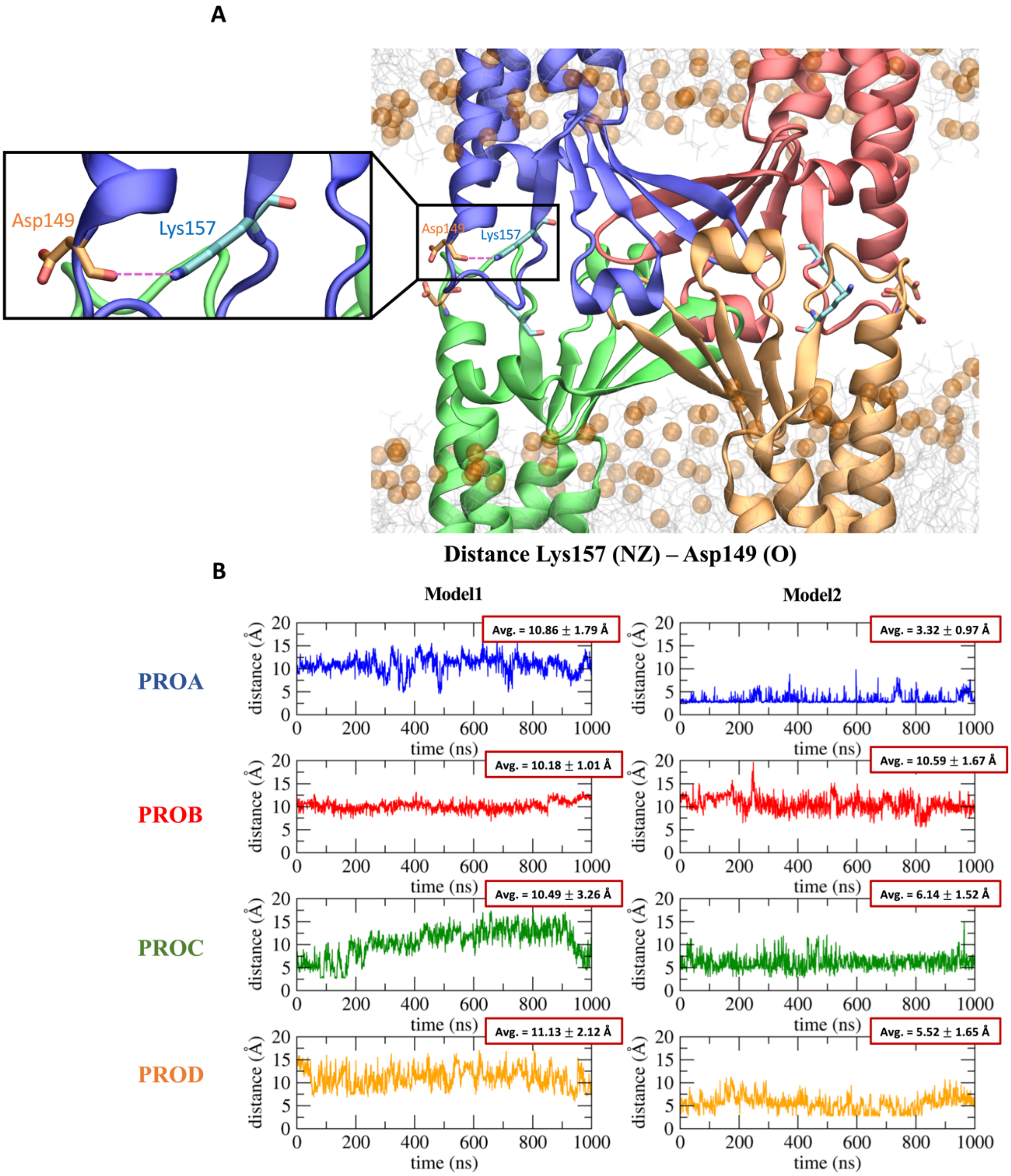


***Figure S5****. Interaction between Lys157 and Asp149 in the four protomers of each model. A) Graphical representation of the two residues at the extremities of the beta-turn in the ECL2. B) Values of distance measured along the trajectory for the individual protomers of each model.*

#### 1.3 Pore Radius

We used the HOLE program[5,6] to calculate the size of the cavity along the pore axis (**Figure S6**). In particular, the narrowest value of the pore radius is localized at the center of the cavity (**Figure S6E,F)**.

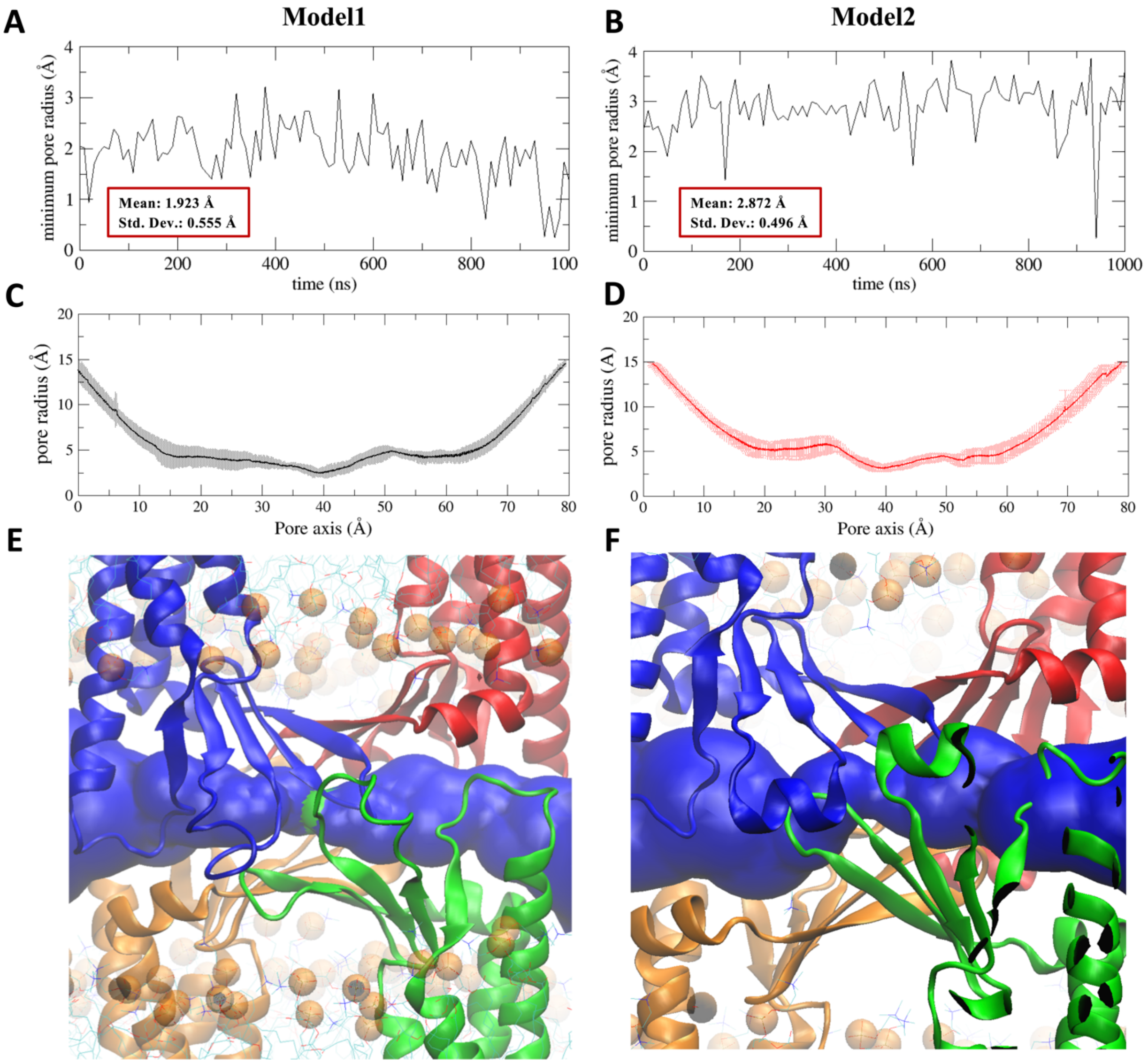


***Figure S6**. Pore radius profiles of Model1 and Model2. A) Minimal pore radius for Model1. B) Minimal pore radius for Model2. C) Average profile along the pore axis for Model1. The grey region identifies the standard deviation. D) Average profile along the pore axis for Model2. The light red region identifies the standard deviation. E) Surface representation of the pore size for Model1. F) Surface representation of the pore size for Model2. The blue-colored areas represent the regions where the radius is at least twice the radius of a single water molecule, while the green-colored section define regions fitting no more than one water molecule.*

In Model2, we observed the preservation of the hydrogen bond between Lys157 and Asp149 for three protomers[4]. Based on this result, we chose Model2 as the best representative structure of the Pore I arrangement, to be used for FE calculations.

#### 2. Block Error Analysis

The FE profiles were calculated with the Umbrella Sampling (US) method[7] in combination with the *weighted histogram analysis method* (WHAM)[8–10], available at http://membrane.urmc.rochester.edu/content/wham. This software implements the bootstrap error analysis to evaluate the uncertainty during the calculation of the FE landscape, which is reported to be underestimated[8,9]. For this reason, we also computed the error estimates using the block analysis, by dividing the trajectories in four blocks[11], as reported in **Figure S7** and **Figure S8**, respectively.

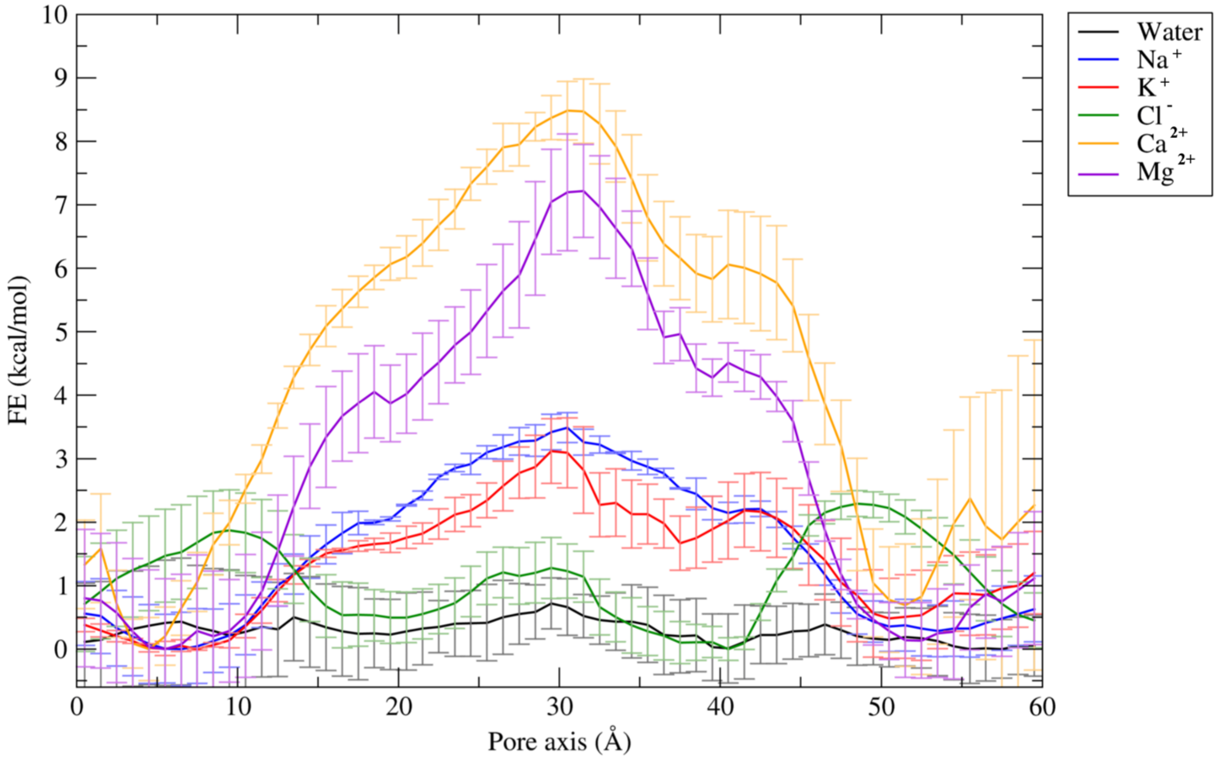


***Figure S7**. Free Energy profiles of water and ions permeating the Pore I. The error, estimated with the block analysis, is represented with bars.*

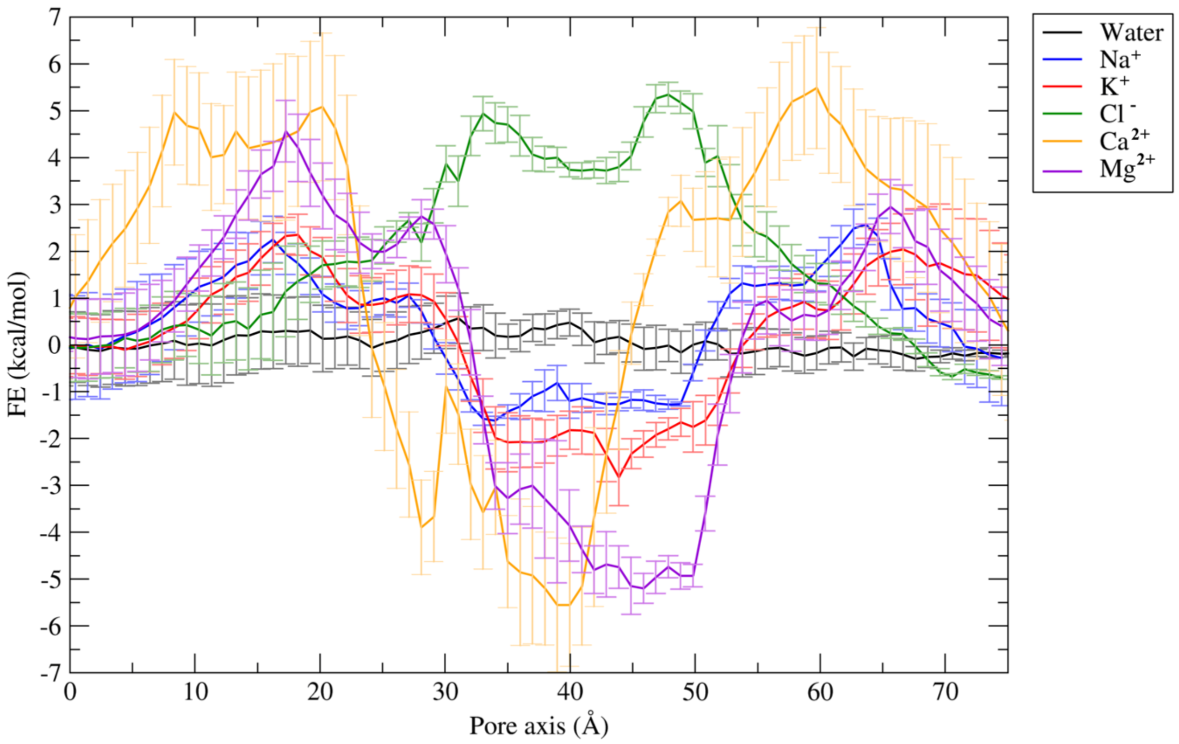


***Figure S8**. Free Energy profiles of water and ions permeating the Pore II. The error, estimated with the block analysis, is represented with bars.*

### 3. Structural Analysis of the Cldn5 monomer

The Cldn5 monomer was homology modelled using the Cldn15 structure as template (PDB ID: 4P79)[2]. In order to assess the structural stability of the model, we performed 110 ns of all-atom MD simulation of the protein embedded in a POPC membrane and solvated with explicit water and counterions. The RMSD of the protein backbone is stationary around 2.25 Å (**Figure S9**). Additionally, we analyzed the network of intramolecular interactions responsible for the stabilization of the tertiary structure[12]. The distances between selected residues are reported in **Table S2** and illustrated in **Figure 10A**. In the inner part of the TM domains, hydrophobic interactions are defined by Trp18, Phe92 and Phe96 of the TM1 and TM2 helices.

In order to assess the conservation of residues involved in the aforementioned interactions , we performed a multiple sequence alignment with Clustal Omega[13,14], using the sequences of the human Cldn5, the mouse Cldn15 (mCldn15, PDB ID: 4P79)[2] and the human Cldn4 (PDB ID: 5B2G)[15]. While the Cldn15 forms barrier to the passage of anions and channels for the cations[11,16], the Cldn4 is mainly supposed to provide an opposite selectivity to the ion permeation[17]. The final alignment is shown in **Figure S10B**. The aromatic Trp30, Trp51, Phe147 and the charged Arg81 residues are conserved among the three homologs, providing an interacting interface between the TM and the ECL domains. Moreover, an analogue cation - $\pi$ was reported in the Cldn15 between the Arg79 and the Phe65 residues[2,12]. In the Cldn4 structure (PDB ID: 5B2G)[15], the electrostatic network formed by the acidic residues Glu48, Asp76 and Asp146 stabilizes the stack conformation of Arg31 and Arg158. Conversely, in the Cldn5 sequence, the two Arg residues are converted in the neutral Asn31 and Tyr158 and the Glu48 negatively charged in Cldn4 becomes the basic Lys48 in the Cldn5 primary sequence. Indeed, the salt bridge (SB) between the Lys48 and the Glu76 was identified, together with the interactions between Lys48 and the Tyr67 and Tyr158 sidechains (**Figure S10A**). In addition, the Gln57 residue of Cldn5 and Cldn4 becomes the acidic Asp55 in the mCldn15, which is pivotal for the cation selectivity of the Cldn15-based TJs[11,16].

Moreover, other key residues in the ECL domain vary among the three Cldns to differentiate the physiological tasks exerted by the proteins[11,16,17].

***Table S2***. *Residue-residue interactions calculated along the 110-ns-long trajectory of the Cldn5 monomer. Distances are calculated between the heavy atoms at the extremities of the sidechains.*

| residues | interaction | Cldn5 region | distance (Å) |
|---|---|---|---|
| **Phe92 -Trp18** | π - π | TM2 - TM1 | 5.33 ± 0.48 |
| **Phe96 - Phe92** | π - π | TM2 - TM2 | 6.49 ± 0.58 |
| **Phe139 - Trp30** | π - π | ECL2 - ECL2 | 6.40 ± 0.31 |
| **Trp30 - Trp51** | π - π | ECL1 - ECL1 | 8.32 ± 0.41 |
| **Arg81 - Trp51** | cation - π | TM2 - ECL1 | 5.70 ± 0.63 |
| **Lys157 – Phe147** | cation - π | ECL2 - TM3 | 5.51 ± 1.48 |
| **Lys48 - Glu76** | SB | ECL1 - TM2 | 6.94 ± 1.59 |
| **Lys157 – Glu159** | SB | ECL2 - ECL2 | 5.79 ± 2.16 |
| **Glu7 - Lys114** | SB | TM1 - TM3 | 7.88 ± 0.45 |
| **Glu44 – Glu57** | HB | ECL1 - ECL1 | 5.87 ± 1.86 |
| **Lys48 - Tyr67** | HB | ECL1 - ECL1 | 4.91 ± 0.90 |
| **Lys48 - Tyr158** | HB | ECL1 - ECL1 | 4.41 ± 0.89 |
| **Asp68 - Arg81** | HB | ECL2 - TM2 | 6.84 ± 1.82 |
| **Phe35 – Gln156** | HB | ECL1 - ECL2 | 3.44 ± 0.51 |

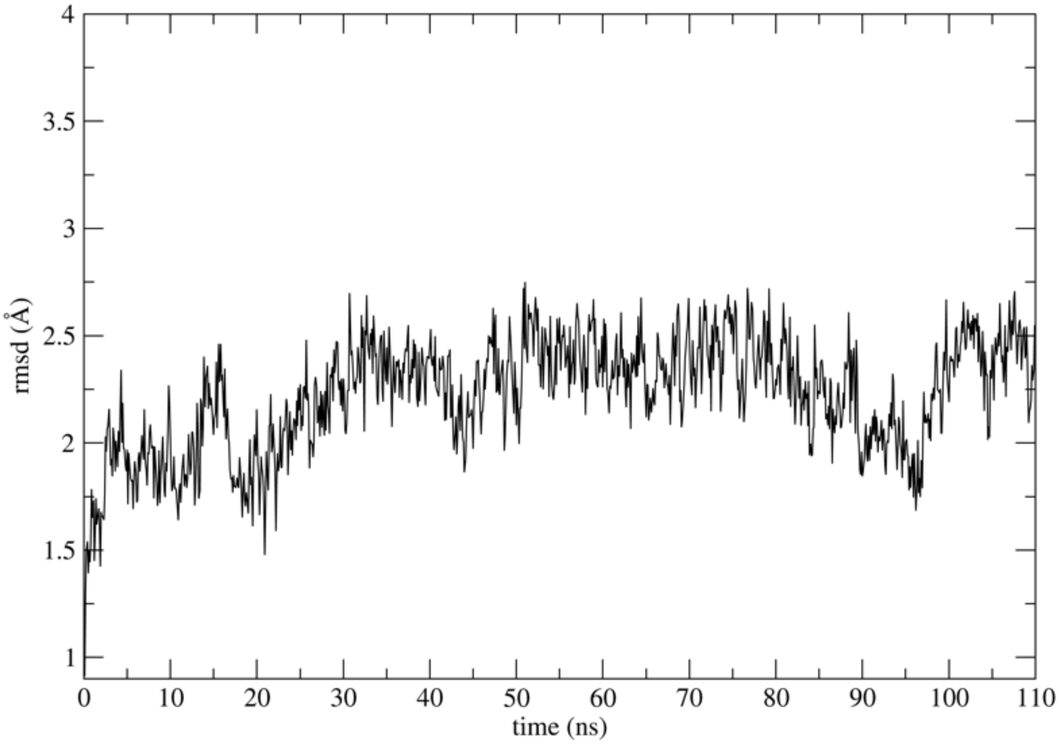


***Figure S9**. RMSD of the Cldn5 backbone.*

**A**

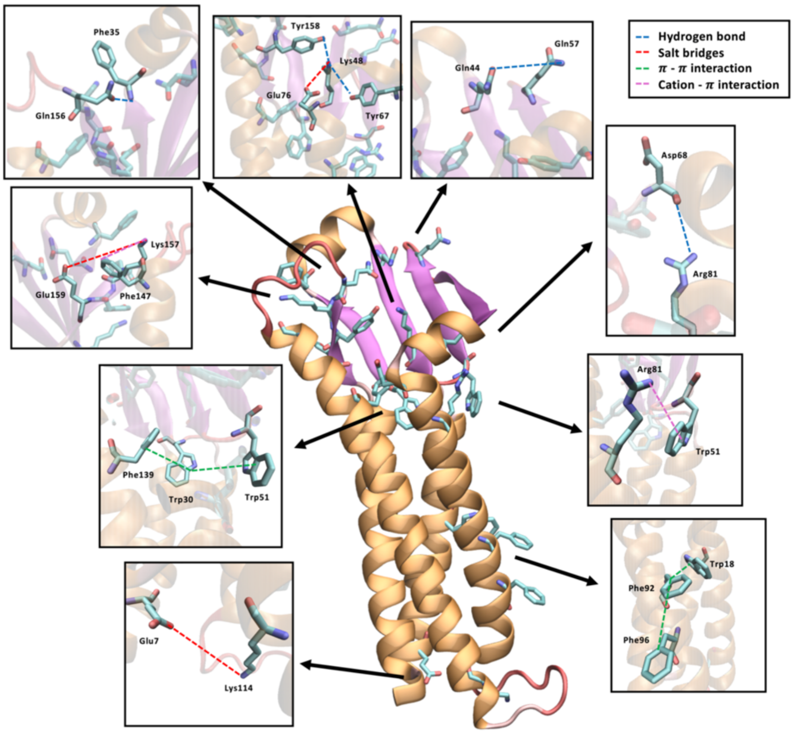


**B**

```
SP|O00501|CLD5_HUMAN  MGSAALEILGLVLCLVGWGGLILACGLPMWQVTAFLDHNIVTAQTTWKGLWMSCVVQSTG 60
SP|Q9Z0S5|CLD15_MOUSE -MSVAVETFGFFMSALGLLMLGLTLSNSYWRVSTV-HGNVITTNTIFENLWYSCATDSLG 58
SP|O14493|CLD4_HUMAN  MASMGLQVMGIALAVLGWLAVMLCCALPMWRVTAFIGSNIVTSQTIWEGLWMNCVVQSTG 60
                        * .:: :*: :. :*   : *  .   *:*::.   *::*::* ::.** .*..:* *

SP|O00501|CLD5_HUMAN  HMQCKVYDSVLALSTEVQAARALTVSAVLLAFVALFVTLAGAQCTTCVAP-GPAKARVAL 119
SP|Q9Z0S5|CLD15_MOUSE VSNCWDFPSMLALSGYVQGCRALMITAILLGFLGLFLGMVGLRCTNVGNMDLSKKAKLLA 118
SP|O14493|CLD4_HUMAN  QMQCKVYDSLLALPQDLQAARALVIISIIVAALGVLLSVVGGKCTNCLED-ESAKAKTMI 119
                        :*  : *:***   :*..*** : ::::. :.::: :.* :**.        **:

SP|O00501|CLD5_HUMAN  TGGVLYLFCGLLALVPLCWFANIVVREFYDPSVPVSQKYELGAALYIGWAATALLMVGGC 179
SP|Q9Z0S5|CLD15_MOUSE IAGTLHILAGACGMVAISWYAVNITTDFFNPLY-AGTKYELGPALYLGWSASLLSILGGI 177
SP|O14493|CLD4_HUMAN  VAGVVFLLAGLMVIVPVSWTAHNIIQDFYNPLVASGQKREMGASLYVGWAASGLLLLGGG 179
                       .*.:.::.*   :* :.* *  :  :*::*    . * *:* :**:**:*: * ::**

SP|O00501|CLD5_HUMAN  LL---CCGAWVCTGRPDLSFPVKYSAPR--------RPTATGDYDKKNYV 218
SP|Q9Z0S5|CLD15_MOUSE CVFSTCCCSSKEEPATRAGLPYKPSTVVIPRATSDESDISFGKYGKNAYV 227
SP|O14493|CLD4_HUMAN  LL---CCN---CPPRTDKPYSAKYSAAR--------SAAASN------YV 209
                       :   **               * *:             : .      **
```

* Fully conserved residues
: High similarity group
· Weak similarity group

***Figure S10****. A) Representation of the Cldn5 structure with enlarged views of the residues involved in the aforementioned interactions. B) Multiple sequence alignment among hCldn5, mCldn15 and hCldn4.*
